# Unveiling the regulatory role of GRP7 in ABA signal-mediated mRNA translation efficiency regulation

**DOI:** 10.1101/2024.01.12.575370

**Authors:** Jing Zhang, Yongxin Xu, Fa’an Tian, Chongsheng He, Xiang Yu, Xiaofei Yang, Yiliang Ding, Jun Xiao

## Abstract

Abscisic acid (ABA) is a crucial phytohormone involved in plant growth and stress responses. Although ABA has been implicated in the regulation of translation efficiency in *Arabidopsis thaliana*, the underlying mechanism remains largely unknown. In this study, we discovered that ABA treatment modulates globally translation efficiency (TE) by affecting pre-rRNA processing in the nucleolus and ribosome distribution status in the cytoplasm. The regulation of TE by ABA was largely abolished in mutants of ABA signaling core components, such as receptors PYRABACTIN RESISTANCE1/PYRABACTIN-LIKE/REGULATORY COMPONENTS OF ABA RECEPTORS (PYR/PYL/RACRs), the protein phosphatase 2Cs (PP2Cs), and the SNF1-related protein kinase 2s (SnRK2s). ABA treatment reduced the protein levels of glycine-rich RNA binding protein 7 (GRP7) in the signaling core components-dependent manner. Ribo-seq and CLIP-seq analyses unveiled GRP7’s role in governing the TE of a substantial proportion of ABA-regulated genes, although independent of directly binding to the respective mRNAs. Furthermore, GRP7 directly bound to pre-rRNA and interacted with Ribosomal protein S6 (RPS6A), RPS14A and RPL36aA in the nucleolus to regulate rRNA processing. Additionally, GRP7 associated with mature polysome in the cytoplasm and is hypersensitive to translation inhibitor anisomycin when loss of function. Collectively, our study unveils the role of GRP7 in mediating translation regulation in ABA signaling, providing a novel regulatory model for plants to response to environmental stresses.

## Introduction

Abscisic acid (ABA) is a crucial phytohormone that regulates plant development and responses to environmental stresses^1,2^. ABA levels in plants are dynamically controlled through tightly regulated biosynthesis and metabolism^2^. The biosynthesis of ABA involves multiple enzymatic steps, with the cleavage of carotenoids by 9-cis-epoxycarotenoid dioxygenase (NCED) enzymes being the rate-limiting step^2^. Under abiotic stress conditions like drought and high salinity, specific signaling pathways are activated, leading to increased ABA accumulation^3,4^. Elevated ABA levels serve as a signal to initiate adaptive responses that enhance stress tolerance, such as stomatal closure and seed dormancy^3,4^. The ABA signaling pathway is initiated by ABA binding to its receptors, PYRABACTIN RESISTANCE1/PYRABACTIN-LIKE/REGULATORY COMPONENTS OF ABA RECEPTORS (PYR/PYL/RACRs), which inhibits protein phosphatase 2C (PP2C) activity. This inhibition releases the suppression of SNF1-related protein kinase 2s (SnRK2s), allowing them to phosphorylate downstream targets like transcription factors and ion channels^5^. Recently, it has been discovered that ABA also plays a significant role in translational regulation in plants^6^. ABA treatment influences ribosome profiling patterns^6^, thereby modulating the expression levels of numerous proteins. However, the precise mechanisms underlying this regulatory process remain largely unknown. Further research is needed to unravel the complex interplay between ABA signaling and translational regulation, deepening our understanding of how ABA controls plant responses to environmental cues.

Ribosomal DNA (rDNA) is transcribed by RNA polymerase I to produce pre-ribosomal RNA (pre-rRNA)^7^. The pre-rRNA undergoes a complex series of processing steps^8^, including cleavage and modification, are carried out by a diverse array of ribonucleoprotein complexes and enzymatic activities present in the nucleolus. There are two pre-rRNA processing pathways coexist in Arabidopsis, in which ITS1 cleavage occurs before or after complete removal of the 5′ ETS, respectively^8,9^. Once the rRNA molecules are processed, they associate with ribosomal proteins to form small (40S) and large (60S) ribosome subunits in the nucleolus^10^. These subunits assemble with transfer RNA and other factors to create functional ribosomes^10,11^. Ribosomes, along with mRNA molecules, form polysomes, allowing multiple ribosomes to translate the same mRNA simultaneously^12^. These polysomes play a crucial role in mediating the efficient and coordinated synthesis of proteins by facilitating the simultaneous translation of multiple ribosomes along the mRNA molecule^13^. This process is essential for protein synthesis, where ribosomes decode mRNA to assemble amino acids into polypeptide chains, generating functional proteins^10^. The formation, assembly, and functioning of ribosomes are crucial for various cellular processes and protein production^10,14^. Alteration of rRNA processing or ribosome biogenesis would influence the translation outcomes.

RNA-binding proteins (RBPs) are conserved proteins found in eukaryotes that interact with RNA molecules to regulate post-transcriptional processes^15,16^. They play diverse roles in RNA metabolism, including RNA processing, alternative splicing, mRNA export, degradation, storage, and translational control^17–20^. RBPs can also act as RNA chaperones, assisting in RNA folding and structure remodeling^21^. Some RBPs are targeted to organelles like chloroplasts or mitochondria and participate in intron splicing, rRNA processing, and translation of plastid mRNAs, crucial for organelle function during stress adaptation^22,23^. RBPs typically contain RNA binding domains (RBDs) responsible for RNA recognition and protein-protein interactions, forming ribonucleoprotein (RNP) complexes^24^. Additionally, RBPs possess auxiliary domains or motifs, such as glycine-rich regions, arginine-rich domains, and serine-arginine repeats, which facilitate protein interactions^24,25^. Among the abiotic stress-associated RBPs, glycine-rich RNA-binding proteins (GR-RBPs) are part of the glycine-rich protein superfamily^26^. GR-RBPs, like GRP7, have been linked to various stresses, including ABA treatment, temperature, drought, and salinity^27–33^. GRP7 exhibits DNA melting and RNase-enhancing activities, potentially preventing the formation of unfavorable RNA secondary structures and facilitating efficient RNA processing, export, and translation under low temperatures^29,30,34^. GRP7 may also act as a shuttle protein, promoting mRNA export from the nucleus to the cytoplasm under cold stress^30^. These findings suggest that GR-RBPs function as RNA chaperones during stress responses. However, their role in translational regulation remains largely unknown.

In this study, we observed significant changes in both pre-rRNA processing in the nucleolus and ribosome status in the cytoplasm following ABA treatment. GR-RBPs, particularly GRP7, play important roles in these changes by directly binding to pre-rRNA and interacting with RPS6A, RPS14A and RPL36aA for proper pre-rRNA processing in the nucleolus. GRP7 also co-fractionated with ribosomes, affecting their activity in translating general mRNAs independent of the directly binding of GRP7.

## Results

### ABA treatment orchestrates dynamic changes in mRNA translation efficiency, pre-rRNA processing, and ribosome distribution

Plant exhibit rapid adjustments in gene expression at both transcriptional and translational levels in response to environmental stress, with the hormone ABA playing a pivotal role^35,36^. We aim to unravel the intricate mechanisms through which ABA treatment influences gene expression dynamics. By conducting RNA-seq and Ribo-seq analyses on 3-day-old Col-0 seedling subjected to ABA or mock treatment, we explored the differential expression of genes and translation efficiency (TE) (Fig. 1a, Fig. S1a).

**Fig. 1.**
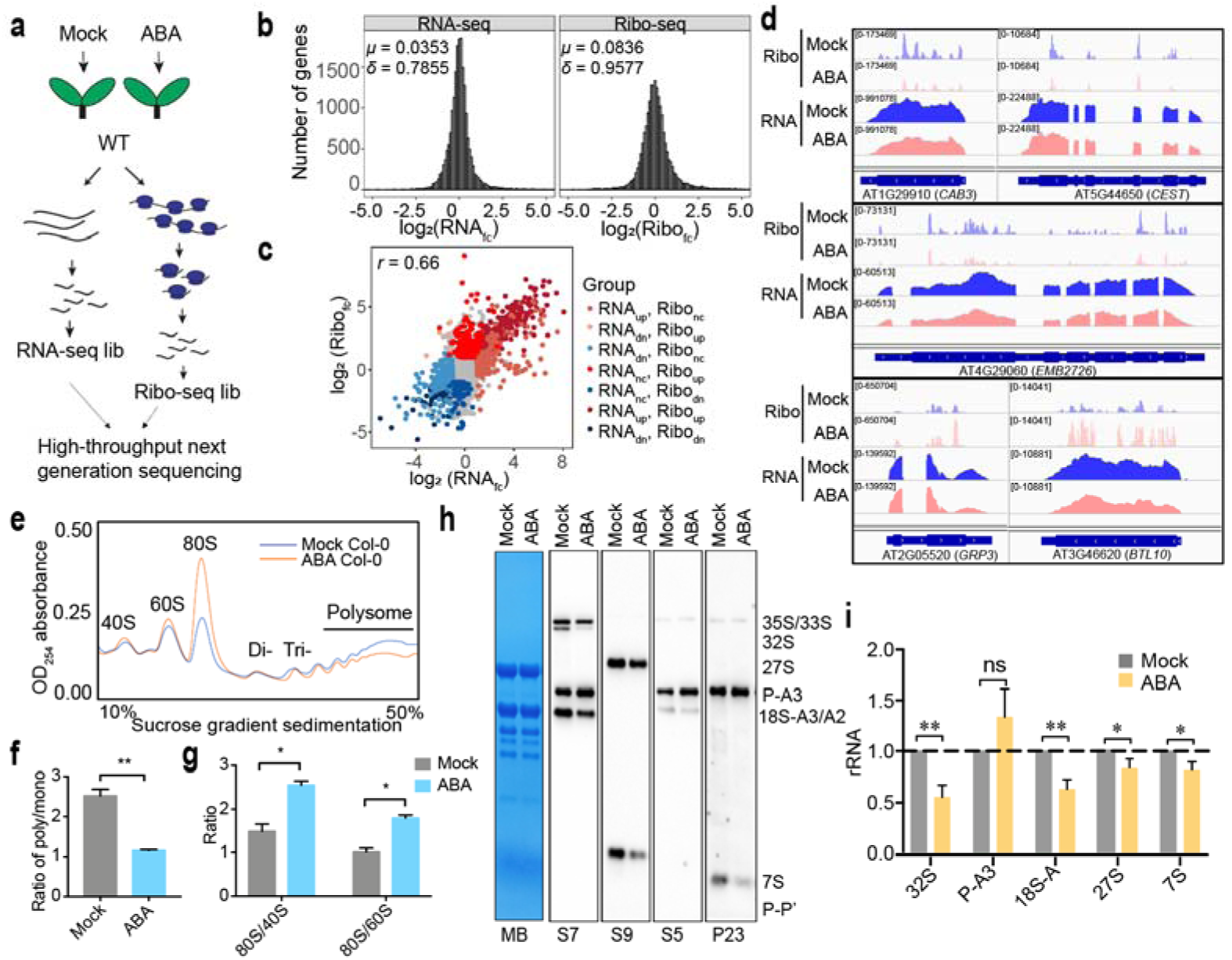
ABA globally modulates translation efficiency via influencing pre-rRNA processing and ribosome status. **a** Schematic representation of RNA-seq and Ribo-seq library construction using Col-0 seedlings with mock or ABA treatment. **b** Histogram illustrating the transcriptional (log_2_(RNA_fc_)) and translational (log_2_(Ribo_fc_)) response to ABA in Col-0. fc, fold change; μ, mean; δ, standard deviation. **c** Relationships between RNAfc and Ribofc; dn, down; nc, no change. **d** IGV screenshots showcasing representative genes with altered RNA-seq and Rio-seq reads under mock and ABA treatment. **e** Polysome profiling revealing the global reduction in TE in Col-0 with ABA treatment. The x-axis indicates the detecting distance on the 10-50% sucrose gradient, ranging from 0 to 75 mm. **f** Quantification of polysome/monosome (poly/mono) ratios for data presented in **(e)**. Student’s *t* test: \*\**p* < 0.01. Two bio-replicates were performed. **g** Ratios of 80S/40S and 80S/60S quantified for the data shown in **(e)**. Student’s *t* test: \*\**p* < 0.01. Two bio-replicates were performed. **h** Northern blot identification depicting the aberrant decrease of 32S, 27S, 18S-A, and 7S rRNAs in Col-0 after ABA treatment. Methylene blue staining (MB) serves as loading control. **i** Quantification of the intensities of different rRNA intermediates. Values were normalized to methylene blue stain for mature 25S and 18S rRNAs and then expressed as a ratio to the intensity observed in Col-0 under mock treatment. The baseline is set to 1, and the standard deviation is indicated as error bar. Student’s *t* test: ns, no significance, \**p* < 0.05; \*\**p* < 0.01. Two bio-replicates were performed.

Under ABA treatment, we identified 5,248 differential expression genes (DEGs) and 1,639 genes with altered TE (Fig. S1b, Supplementary Dataset 1 and 2). Notably, the range of the fold changes (ABA/Mock) in translation (Ribo_fc_) exceeded that of the fold changes (ABA/Mock) in transcription (RNA_fc_), highlighting the dynamic nature of translation compared to transcription under ABA treatment (Fig. 1b). While a strong correlation existed between transcription and translation levels within the same sample (Fig. S1c), the ABA/mock treatment induced a notable shift (Fig. 1c), leading to a poor correction in fold changes between transcription and translation, indicating that ABA treatment regulates global TE. Among genes exhibiting changes in TE, there is an enrichment of gene ontology (GO) terms related to photosynthesis, chloroplast organization, and translation in down-regulated TE genes. Conversely, up-regulated TE genes are enriched in GO terms associated with response to abscisic acid, water deprivation, and cold (Fig. S1d, e). This is consistent with the activation of defense processes while the shutdown of normal growth occurs under ABA treatment. Notably, this includes photosynthesis and chloroplast biogenesis genes, such as Chlorophyll A/B binding protein 3 (*CAB3*, *AT1G29910*)^37^, Chloroplast protein-enhancing stress tolerance (*CEST*, *AT5G44650*)^38^, Embryo defective 2726 (*EMB2726*, *AT4G29060*)^39^, and genes function in plant stress response, such as Glycine-rich protein 3 (*GRP3*, *AT2G05520*)^40^, and BCA2A ZINC FINGER ATL 10 (*BTL10*, *AT3G46620*)^41^ (Fig. 1d), emphasizing the pivotal role of translation regulation in the response to ABA.

We performed polysome profiling on Col-0 seedlings to assess the global impact of ABA treatment on translation. Notably, ABA treatment resulted in a significant reduction in RNA content in the polyribosome fractions compared to the mock treatment. The polysome/monosome ratio decreased, indicating a global inhibition of translation by ABA (Fig. 1e, f), as reported before^6^. Furthermore, ABA treatment increased the ratios of 80S/60S and 80S/40S compared to the mock treatment, indicating potential effects on ribosome assembly *in vivo* (Fig. 1g). Subsequently, we explored the impact of ABA on pre-rRNA processing, a critical step in ribosome biogenesis^10^. *Arabidopsis* employs two pre-rRNA processing pathways, with cleavage at the A3 site occurring either before or after complete removal of the 5’ETS at the P site (Fig. S2a). Northern blot analyses revealed abnormal decreases in 32S, 18S-A2 and/or 18S-A3, 27S, and 7S rRNA intermediates under ABA treatment in Col-0 (Fig. 1h, i). Intriguingly, 32S rRNA is associated with the minor alternative pre-rRNA processing pathway. Consequently, ABA treatment disrupts processing intermediates from the 5’ETS first pathway, leading to the inhibition of the minor pre-rRNA processing pathway (Fig. S2b).

### ABA signaling pathway components regulate global mRNA translation efficiency

We examined the impact of ABA biosynthesis and signaling pathway factors on the regulation of mRNA TE. Specifically, we investigated the *nced6* mutants^2^, which affects the speed-limiting ABA biosynthesis enzyme, the ABA receptors mutant *pyr1pyl1/2/4*^42^, the negative regulator of ABA signaling key phosphatase mutant *pp2c 3m*^5^, and the key kinase mutant *snrk2.2/2.3/2.6*^43^.

In our polysome profiling analysis, the *pp2c 3m*, mutation of the negative regulator for ABA signaling, exhibited a reduction in the polysome/monosome ratio and an increase in 80S/40S and 80S/60S compared to Col-0 under mock conditions, similar as Col-0 with ABA treatment (Fig. 2a, b, c, Fig. S3). Whereas, in *pyr1pyl1/2/4* mutant, these indexes don’t change much compared to Col-0. Mutation of core kinase SnRK2s also displayed reduced polysome/monosome ratio and increased 80S/40S and 80S/60S ratios, likely due to the increased 80S monosome solely in *snrk2.2/2.3/2.6* compared to Col-0 under mock condition (Fig. 2a, b, c, Fig. S3).

**Fig. 2.**
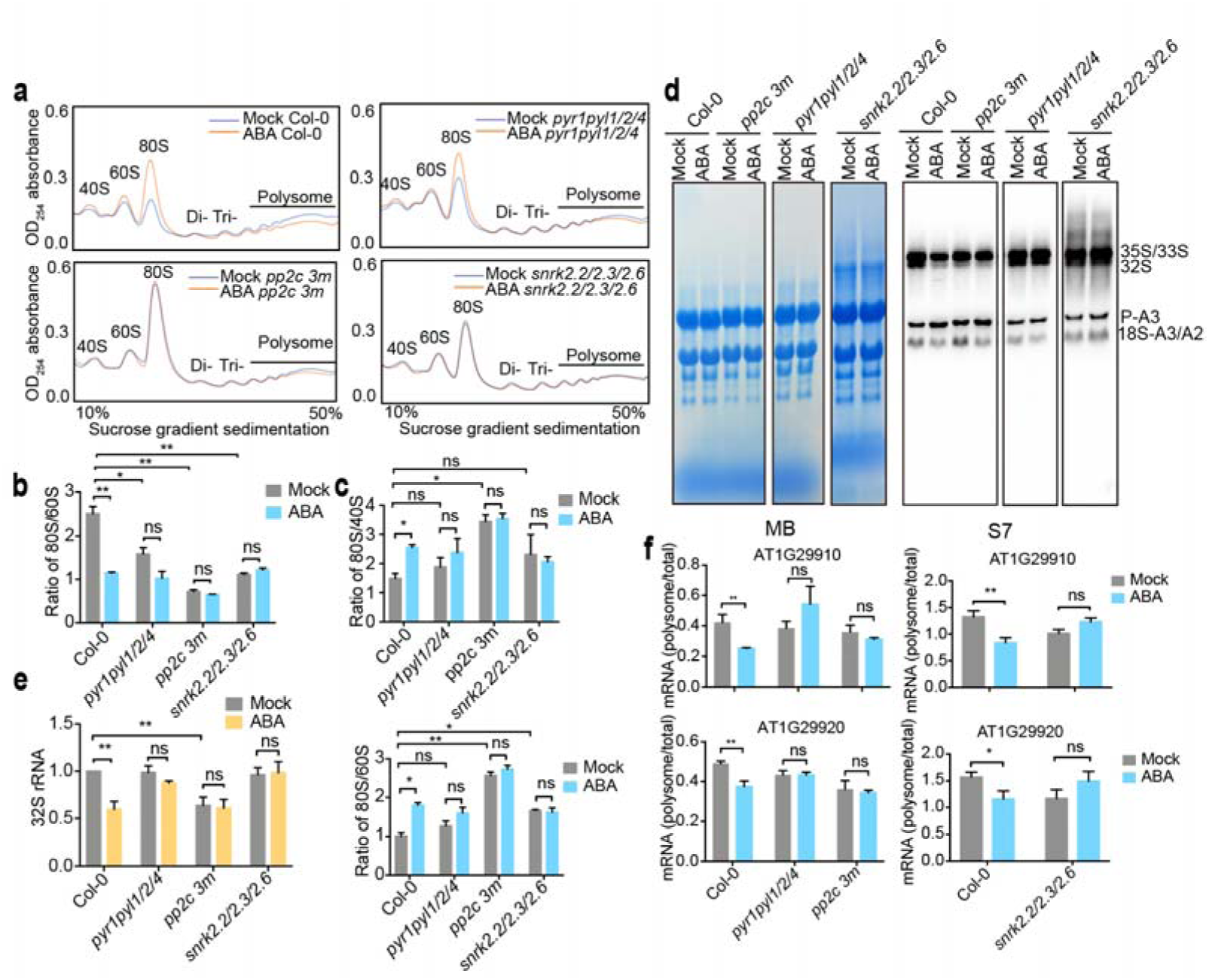
ABA affects translation efficiency through its signaling pathway factors. **a** Polysome profiling was analyzed with sucrose gradient sedimentation from 10-50% for Col-0, *pyr1/pyl1/2/4*, *pp2c 3m* and *snrk2.2/2.3/2.6* with mock and ABA treatment. **b** Quantification of polysome/monosome (poly/mono) ratios for data presented in **(a)**. Student’s *t* test: ns, no significance, \**p* < 0.05; \*\**p* < 0.01. Two to three bio-replicates were performed. **c** Ratios of 80S/40S and 80S/60S quantified for the data shown in **(a)**. Student’s *t* test: ns, no significance, \**p* < 0.05; \*\**p* < 0.01. Two bio-replicates were performed. **d** Northern blot identification depicting the aberrant pre-rRNA processing in Col-0, *pyr1/pyl1/2/4*, *pp2c 3m* and *snrk2.2/2.3/2.6* after ABA treatment. Methylene blue staining (MB) serves as loading control. **e** Quantification of the intensities of 32S rRNA in Col-0, *pyr1/pyl1/2/4*, *pp2c 3m* and *snrk2.2/2.3/2.6* after ABA treatment. Values were normalized to methylene blue stain for mature 25S and 18S rRNAs and then expressed as a ratio to the intensity observed in Col-0 with mock. The baseline is set to 1, and the standard deviation is indicated as error bar. Student’s *t* test: ns, no significance, \**p* < 0.05; \*\**p* < 0.01. Two bio-replicates were performed. **f** qRT-PCR was performed to quantified the translation efficiency of several genes upon ABA treatment in Col-0 and different mutants. Student’s *t* test: ns, no significance, \**p* < 0.05; \*\**p* < 0.01. Two bio-replicates were performed.

Thus, alteration of ABA signaling could influence polysome profiling. Moreover, ABA treatment reduced ribosome loading efficiency and increase 80S/40S and 80S/60S ratios in the *nced6*, comparable to Col-0 (Fig. S4a, b, c). However, in *pyr1pyl1/2/4*, *pp2c 3m* and *snrk2.2/2.3/2.6* mutants, the impact of ABA treatment on reducing polysome loading (Fig. 2a, b) and increasing 80S/40S and 80S/40S ratios was notably diminished compared to Col-0 (Fig. 2a, c). This further confirmed that ABA signaling core components are required in regulating polysome profile.

We extended our analysis to pre-rRNA processing in these mutants. ABA treatment in the *nced6* mutant suppressed the accumulation of 32S rRNA intermediates and inhibited the minor pre-rRNA processing pathway, similar to Col-0 (Fig. S4 d, e). Conversely, these effects were abolished in the *pyr1pyl1/2/4, pp2c 3m* and *snrk2.2/2.3/2.6* mutants, highlighting the essential role of ABA perception and transduction through key phosphatase and kinase in this regulation (Fig. 2d, e). Under mock conditions, the *pp2c 3m* mutant exhibits a decrease in 32S rRNA, similar to the trend observed in Col-0 under ABA treatment (Fig. 2d, e). We further quantified the TE of several photosynthesis related genes upon ABA treatment in different mutants. While Col-0 showed reduction TE of *CAB3* (*At1g29910*) and *CAB2* (*At1g29920*) in response to ABA, *pyr1pyl1/2/4*, *pp2c 3m*, and *snrk2.2/2.3/2.6* mutants did not exhibit significant changes (Fig. 2f).

In summary, the ABA-mediated regulation of gene translation primarily relies on the signaling transduction from receptors to signaling components including core phosphatase and kinase.

### ABA signaling reduces GRP7 protein level to regulate translation efficiency

To decipher the impact of ABA signaling on translation regulation, we established specific criteria to identify potential mediators, focusing on factors regulated by ABA signaling, involved in ABA-regulated plant development, and whose mutations affect translation efficiency.

GRPs, a class of RNA binding proteins highly expressed in plant^44^, emerged as promising candidates. ABA treatment led to a significant reduction in the protein level of GRP7, as demonstrated by western blot analysis with anti-GRP7 antibody^45^ (Fig. 3a), which was not attributed to protein stability changes but rather to the translational efficiency reduction of GRP7 mRNA (Fig. S5a-c). Importantly, the reduction of GRP7 protein levels upon ABA treatment was abolished in signaling component mutants, such as *pyr1pyl1/2/4, pp2c 3m* and *snrk2.2/2.3/2.6* (Fig. 3a), while the ABA biosynthesis mutant *nced6* exhibited a clear decline in GRP7 protein level under ABA treatment (Fig. S5d).

**Fig. 3.**
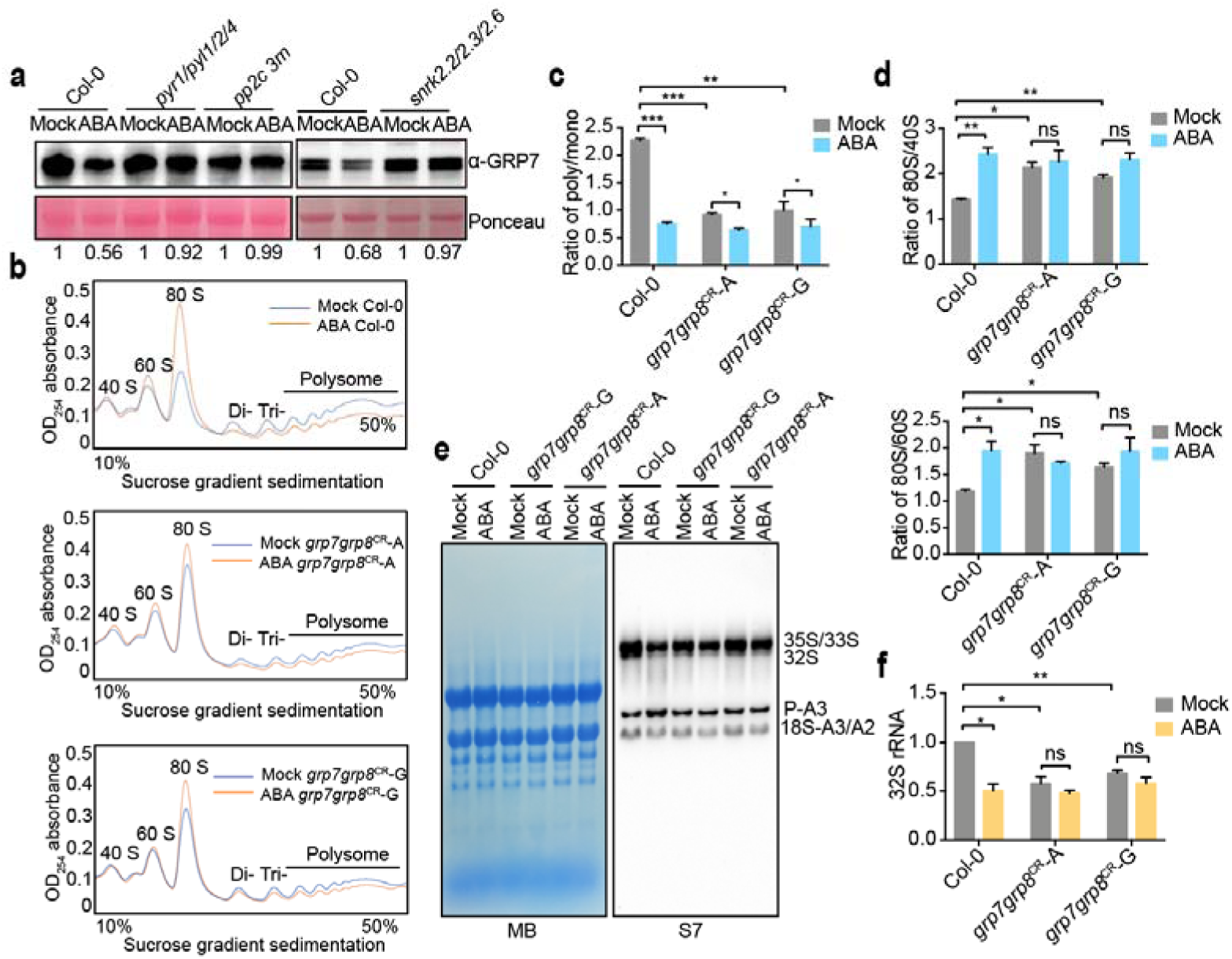
ABA signal reduces GRP7 protein to participate in translation regulation. **a** Detection of GRP7 protein levels in Col-0, *pyr1pyl1/2/4*, *pp2c 3m* and *snrk2.2/2.3/2.6* following ABA treatment using an anti-GRP7 antibody. **b** Polysome profiles of Col-0 and *grp7grp8*^CR^-A/G under mock and ABA conditions. **c** Quantification of polysome/monosome (poly/mono) ratios for data presented in **(b).** Student’s *t* test: \**p* < 0.05; \*\**p* < 0.01; \*\*\**p* < 0.001. Two bio-replicates were performed. **d** Ratios of 80S/40S and 80S/60S quantified for the data shown in **(b)**. Student’s *t* test: \*\**p* < 0.01. Two bio-replicates were performed. **e** Northern blot identification depicting the aberrant pre-rRNA processing in Col-0 and *grp7grp8*^CR^-A/G with mock and ABA treatment. Methylene blue staining (MB) serves as loading control. **f** Quantification of the intensities of 32S rRNA in Col-0 and *grp7grp8*^CR^-A/G with mono or ABA treatment. Student’s *t* test: \**p* < 0.05; \*\**p* < 0.01; ns, no significance. More than two bio-replicates were performed.

GRP8 closely resembles GRP7, displaying high sequence similarity and functional redundancy in impacting alternative splicing^46^ and flowering^47^. To further elucidate their roles in ABA-regulated plant development and potential involvement in mRNA TE regulation, we generated a *grp7grp8* double mutant through genome editing of GRP8 in *grp7-1* background (Fig. S5e). A single insertion of nucleotide A or G into the first exon of *GRP8* led to pre-mature termination at either eighth or 36th amino acid position (Fig. S5e). Previous studies demonstrated that the *grp7-1* mutant confers hypersensitivity to ABA during seed germination process^27,33^. Notably, both *grp7grp8^CR^*-A/G mutant lines exhibited significantly higher ABA sensitivity than the *grp7-1* or *grp8^CR^*-A/G single mutants (Fig. S5f, g), indicating the redundant functions of GRP7 and GRP8 in ABA-mediated inhibition of cotyledon greening. We further analyzed the polysome profiles of Col-0 and *grp7grp8^CR^*-A/G mutant lines with mock and ABA treatment (Fig. 3b). Remarkably, both *grp7grp8^CR^*-A/G lines exhibited reduced ribosome loading efficiency compared to Col-0 at mock condition, similar to the effect of ABA treatment in Col-0 but with a lesser reduction (Fig. 3c, Fig. S5h, i). Importantly, ABA treatment still resulted in reduced ribosome loading efficiency in both *grp7grp8*^CR^ -A/G lines (Fig. 3c), suggesting that GRP7&8 partially mediate the influence of ABA on ribosome profile regulation. Additionally, compared with Col-0, the *grp7grp8*^CR^-A/G lines showed increased ratios of 80S/60S and 80S/40S under mock condition (Fig. 3d). Notably, ABA treatment did not further increase ratios of 80S/60S and 80S/40S in *grp7grp8*^CR^-A/G lines compare with mock (Fig. 3d), indicating that GRP7&8 mediate the major influence of ABA on ribosome assembly *in vivo*.

We further investigated the potential involvement of GRP7&8 in the regulation of the minor pre-rRNA processing pathway, which is inhibited by ABA treatment. Northern blot analysis revealed a decrease in 32S levels in *grp7grp8^CR^*-A/G lines under mock treatment compared to Col-0, similar to the effect of ABA treatment in Col-0 (Fig. 3e, f). Furthermore, ABA treatment did not further affect the 32S intermediate in *grp7grp8^CR^*-A/G lines (Fig. 3e, f), indicating that GRP7&8 could mediate the ABA-induced regulation of pre-rRNA processing.

### GRP7&8 mediate ABA signaling-regulated mRNA translation efficiency

To assess the role of GRP7&8 in translation regulation, we performed Ribo-seq and RNA-seq analysis on Col-0 and *grp7grp8^CR^*-A. The range of the fold changes (*grp7grp8^CR^*-A/Col-0) in translation (Ribo_fc_) exceeded that of the fold changes (*grp7grp8^CR^*-A/Col-0) in transcription (RNA_fc_), underscoring the involvement of GRP7&8 in translation regulation (Fig. 4a, Supplementary Dataset 3, 4). The correlation observed between the fold changes of transcription and translation for *grp7grp8^CR^*-A/Col-0 (Fig. 4b) suggests that the fold changes of mRNA tend to underestimate the fold changes of protein^48,49^, indicating GRP7&8 regulates translation. Among the genes with TE changes, GO terms related to photosynthesis, chloroplast thylakoid, and ribosome were enriched among TE down-regulated genes, while GO terms associated with the response to abscisic acid and water deprivation were enriched among TE up-regulated genes (Fig. S6a). This TE alteration pattern mirrors that observed in Col-0 upon ABA treatment (Fig. S1e). Remarkably, genes exhibiting TE reduction upon ABA treatment showed a significant overlap with TE reduction in *grp7grp8^CR^*-A compared to Col-0 (Fig. 4c).

**Fig. 4.**
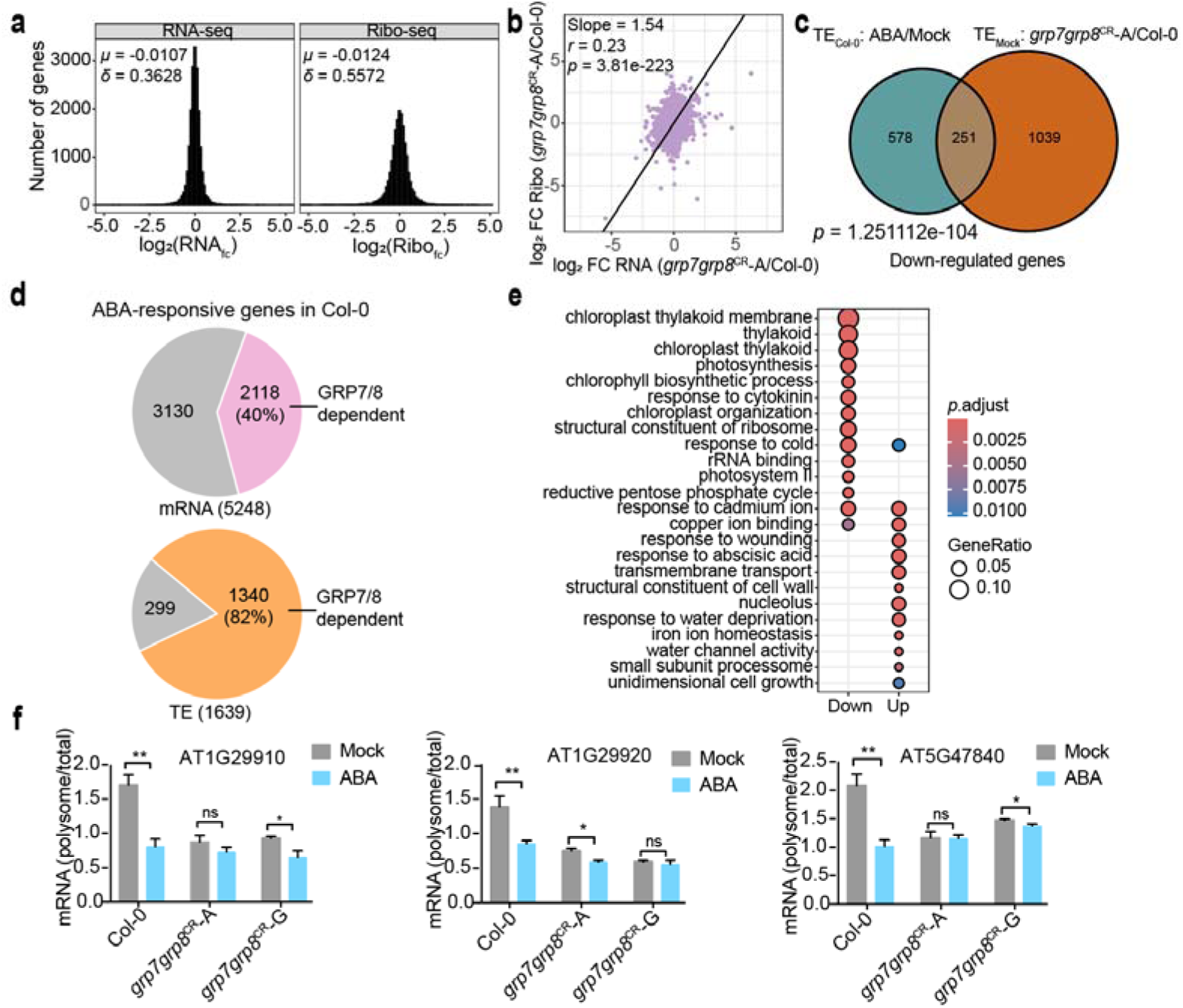
Comparative analysis of transcription and translation alteration in *grp7grp8* -A and Col-0 under ABA treatment. **a** Histogram depicting transcription (log2(RNAfc)) and translation (log2(Ribofc)) changes in *grp7grp8*^CR^-A. fc, fold change; μ, mean; δ, standard deviation. **b** Relationships between RNAfc and Ribofc in Col-0 and *grp7grp8*^CR^-A. Slope = 1.54. **c** Venn diagram illustrating the number of genes co-downregulated by ABA treatment and *grp7grp8*^CR^-A mutation. **d** Comparison of the ABA responsive-genes at mRNA and TE levels in Col-0 and *grp7grp8*^CR^-A. Significantly differentially expressed genes in Col-0 were compared with those in *grp7grp8*^CR^-A using Venn diagrams. **e** GO enrichment analysis of genes downregulated and upregulated in TE of Col-0 dependent on GRP7&8. The size of points indicates the gene ratio in each GO category, and the color scale represents *p*.adjust value. **f** Validation of representative genes. qRT-PCR measuring transcript levels in total RNA and polysome RNA of Col-0 and *grp7grp8*^CR^-A. Detection of photosynthesis genes (*AT1G29910* and *AT1G29920*) and the energy homeostasis gene (*AT5G47840*). *PP2A* was used as an internal reference. Student’s *t*-test: \**p* < 0.05; \*\**p* < 0.01. Two bio-replicates were performed.

To explore whether GRP7&8 mediate mRNA TE regulation at the whole-genome level during ABA treatment, we compared Ribo-seq and RNA-seq data from Col-0 and *grp7grp8*^CR^*-A* under ABA treatment. Quality assessment revealed a high degree of reproducibility between replicates (Fig. S6b-d, Supplementary Data5). We performed a comprehensive gene expression analysis in *grp7grp8*^CR^*-A* mutant compared to Col-0. Among the 5,248 ABA-induced differentially expressed genes in Col-0, the transcriptional response of 2,118 (∼40%) genes was blocked by *grp7grp8*^CR^*-A* mutation (Fig. 4e, Supplementary Data 1, 6, for detailed data process, see method). Among the 1,639 TE-altered genes in Col-0, 1,340 (∼82%) genes were disrupted in *grp7grp8*^CR^*-A* mutant (Fig. 4d, Supplementary Data 2, 7). Notably, among those GRP7&8-dependent and ABA-regulated altered TE genes, photosynthesis and chloroplast organization-related genes were down-regulated, while abiotic stress response genes were up-regulated (Fig. 4e). We further validated the down-regulation of photosynthesis genes *CAB3* (*At1g29910*) and *CAB2* (*At1g29920*) and energy homeostasis gene Adenosine monophosphate kinase (AMK2, *AT5G47840*) by Polysome RNA qRT-PCR^50^, confirming their down-regulation after ABA treatment in a GRP7&8-dependent manner (Fig. 4f). This comprehensive analysis establishes GRP7&8 as key mediators of the ABA-regulated translational response.

### GRP7 binds to pre-rRNA and interacts with ribosomal proteins in the nucleolus

GRP7, a nucleocytoplasmic shuttling protein, exhibits a dual localization in the cytoplasm and nucleus^45^, with a notable presence in the nucleolus (Fig. 5a), as shown co-localization with the nucleolus marker AtFIB2-mCherry in tobacco leaves^51^. This unique sub-nuclear distribution prompted an exploration of GRP7’s potential involvement in rRNA process.

**Fig. 5.**
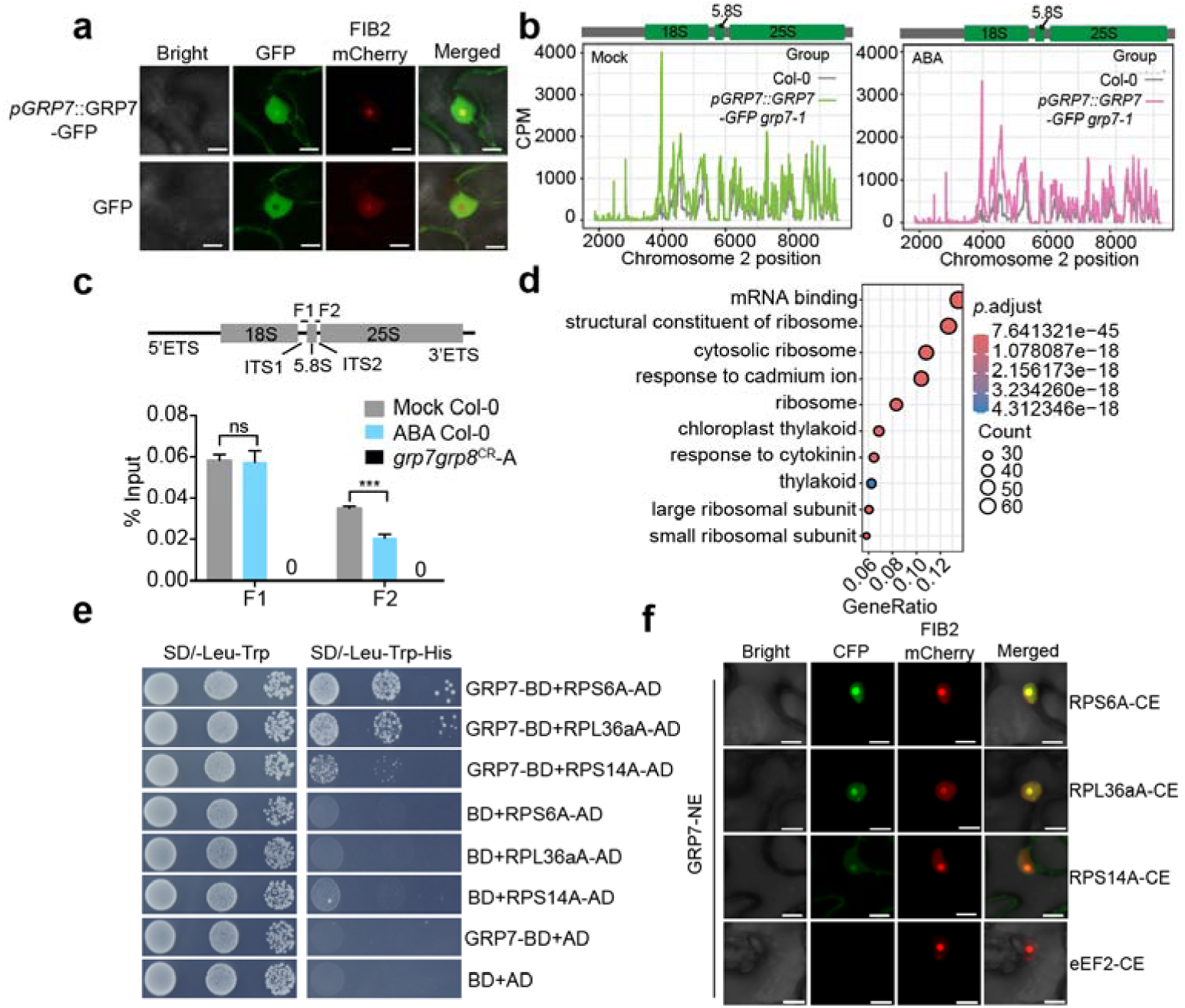
GRP7 binds to pre-rRNA and interacts with ribosomal proteins in nucleolus. **a** Subcellular localization of GRP7 protein. A plasmid expression GRP7-GFP fusion protein was transformed into tobacco along with FIB2-mCherry, a marker for nucleolus. GFP alone was used as the control. Scale bars, 10 µm. **b** Read density of GRP7 binding to rRNA on chromosome 2 under mock (green) and ABA (pink) conditions. Gray lines represent GRP7 binding levels in corresponding control samples. **c** RIP-qPCR detected the association between GRP7 and pre-rRNAs in Col-0 after ABA treatment. The upper diagram indicates the locations of specific fragments (F1 and F2) amplified by qPCR with specific primers. *grp7grp8*^CR^-A was used as the negative control. Student’s *t*-test: \*\*\**p* < 0.001; ns, not significant. Two bio-replicates were performed. **d** GO enrichment of proteins associated with GRP7 under mock (pink) and ABA (green) condition by IP-MS. The x-axis indicates the fold enrichment. **e** Yeast-two-hybrid assay for the interaction between GRP7 and ribosome biogenesis factors. **f** BiFC assay for the interaction between GRP7 and ribosome biogenesis factors in tobacco leaves. FIB2-mCherry serves as a nucleolar maker. eEF2 is the negative control. Scale bar, 10 µm.

Having established GRP7’s localisation, we delved into its role in pre-rRNA processing. Previous studies highlighted GRP7’s direct binding to mRNA, influencing mRNA stability and alternative splicing^46,52^. To extend this understanding, we performed protein-RNA crosslinking and immunoprecipitation (CLIP)-seq analysis. Using the *pGRP7::GRP7-GFP grp7-1* complementation line^53^ and Col-0, coupled with GFP-Trap^54^, we identified 18S and 25S pre-rRNA as robust targets of GRP7, both under mock and ABA conditions (Fig. S7a. b, Supplementary Data 8). The precipitated rRNA, while slightly reduced in intensity under ABA might be due to reduced GRP7 protein level upon ABA treatment, suggests GRP7’s direct and dynamic involvement in pre-rRNA processing (Fig. 5b, Fig. S7c). Complementary RNA immunoprecipitation (RIP) experiments using GRP7 antibody and nuclear fractions of Col-0 seedlings further affirmed specific enrichment of pre-rRNA fragments in the Col-0 samples^55^. Remarkably, this interaction was significantly diminished in *grp7grp8*^CR^*-A* (Fig. 5c), serving as a negative control. Intriguingly, ABA treatment exhibited a modest impact on the GRP7-pre-rRNA binding (Fig. 5c).

Following rRNA processing, ribosomal proteins associate with rRNA to form small (40S) and large (60S) ribosome subunits^10^. We further conducted GFP-tag immunoprecipitation followed by mass spectrometry (IP-MS) analysis using *pGRP7:GRP7-GFP grp7-1* line, with Col-0 as control (Fig. S7d). Proteins identified in the complementation line but not Col-0 in at least two out of three replicates were considered as putative GRP7-interacting proteins (Supplementary Data 9). Interestingly, GRP7 was association with a spectrum of ribosomal proteins from both the large (60S) and small (40S) ribosomal subunits (Fig. 5d). This interaction was further confirmed through yeast two-hybrid (Y2H) assays, validating direct associations between GRP7 and RPS6A, RPS14A, and RPL36A, key components of the large and small ribosomal subunits (Fig. 5e). The bimolecular fluorescence complementation (BiFC) assay further substantiated these interactions, primarily locating them within the nucleolus (Fig. 5f).

These findings underscore the multifaceted role of GRP7 in the intricate landscape of nucleolar pre-rRNA processing.

### GRP7 affects mRNA translation process via association with mature ribosomes in the cytoplasm

To unravel the potential impact of GRP7 on the mRNA translation process in the cytoplasm, we conducted sucrose gradient centrifugation of *pACTIN2::FLAG-RPL18* lysates to probe the association of GRP7 with mature ribosomes (Fig. 6a). The expected enrichment of RPL18 in the 60S, 80S monosome and polysome fractions, and RPS6 mainly in the 40S, 80S monosome and polysome fractions validated the procedure. Intriguingly, GRP7 co-fractionated with the 40S, 60S, monosome, and polysome fractions, suggesting its active participation in the translation process. To rule out the possibility of non-ribosomal complex co-fractionation, we treated lysates with EDTA to disassemble 80S monosomes into 40S and 60S ribosomal subunits, resulting in the depletion of GRP7 from monosome and polysome fractions (Fig. S8). Additionally, RNase A treatment, inducing the accumulation of 80S monosomes, led to a substantial reduction in GRP7 levels in polysome fractions (Fig. S8). These findings robustly support the direct interaction between GRP7 and actively translating mature ribosomes.

**Fig. 6.**
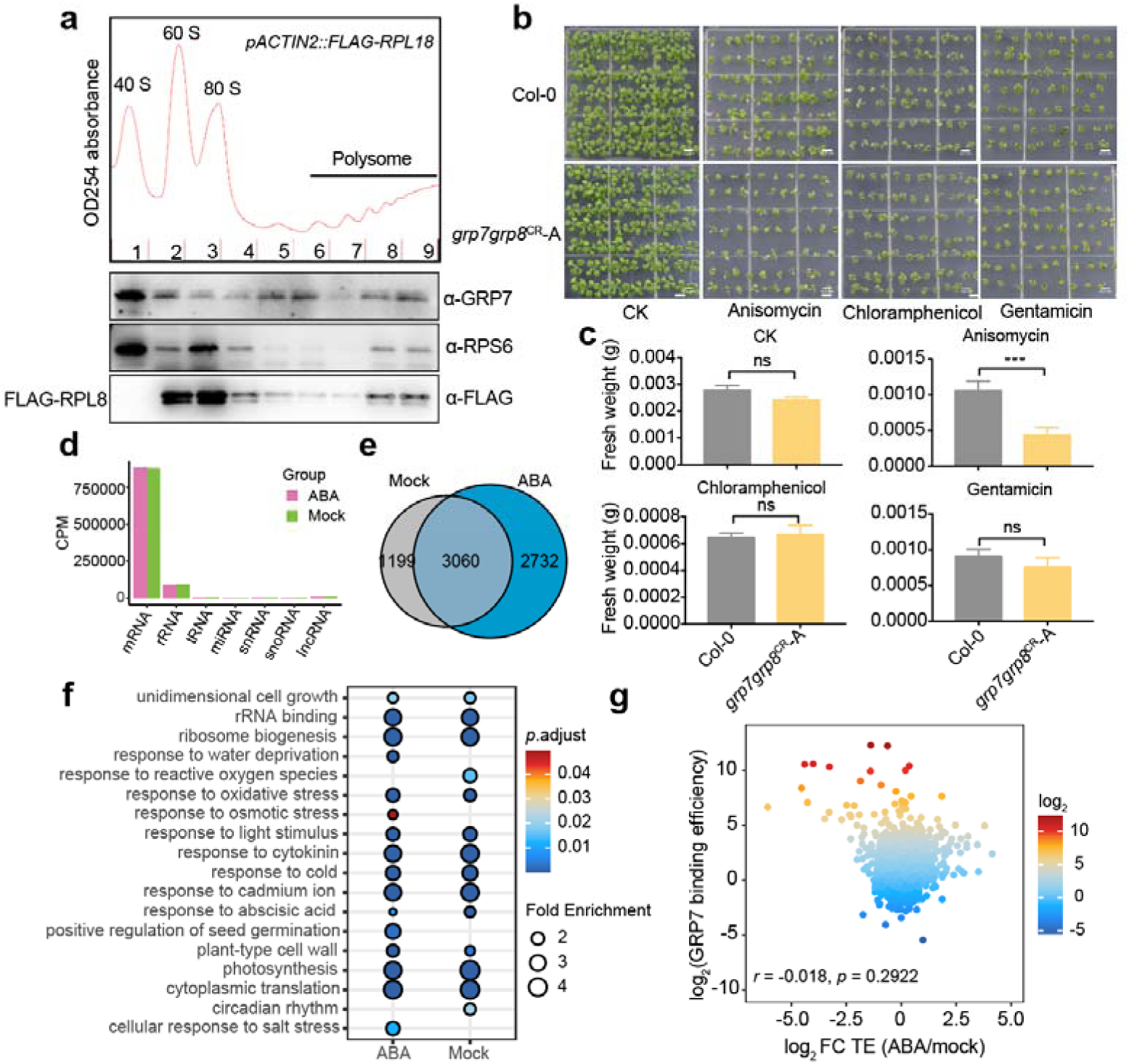
GRP7 regulates mRNA translation in cytoplasm. **a** Polysome profile and western blot analysis of GRP7 associated with ribosome. Top, absorbance plot of *pACTIN2::FLAG-RPL18* fractionated through a sucrose gradient. Bottom, western blots of GRP7, RPS6 and FLAG-RPL18 from corresponding fractions. **b** Antibiotic treatment assay. Col-0 and *grp7grp8*^CR^-A seedlings were grown on 1/2 MS medium containing 0.5 μM anisomycin, 5 μg/mL chloramphenicol, and 40 μg/mL gentamicin. Scale bars, 2 mm. **c** Fresh weight of the Col-0 and *grp7grp8*^CR^-A grown in 1/2 MS medium supplemented with different antibiotics. n > 30 for each sample. The *p-*values were calculated using Student’s *t* test: \*\*\**p* < 0.001; ns, not significant. Three bio-replicates were performed. **d** Bar chart showing fractions of GRP7 CLIP reads unambiguously mapping to mRNAs, rRNAs and other types of RNA. **e** Venn diagram illustrating the intersection of genes bound by GRP7 under mock and ABA treatment. **f** GO enrichment of GRP7 binding genes in mock and ABA treatment conditions. **g** Correlation analysis between GRP7 target gene binding strength and the fold change in translation efficiency under ABA treatment.

To assess the impact of GRP7 on ribosomal functions, we examined the sensitivity of the *grp7grp8*^CR^*-A* mutant to various protein translation inhibitors. Notably, the mutant displayed heightened sensitivity to anisomycin (Fig. 6b, c), an inhibitor of peptide bond formation by competing with amino acids for access to the peptidyltransferase center (A-site)^6^, consistent with a role for GRP7 in protein translation.

Given GRP7’s capacity to bind mRNA (Fig. 6d), we explored whether its regulation of mRNA TE correlated with direct target mRNA. Analyzing CLIP-seq data from *pGRP7::GRP7-GFP grp7-1* revealed significant overlap in mRNA binding, both under mock and ABA-treated conditions (Fig. 6e). Enrichment of GO terms related to abiotic stress, translation, and photosynthesis among GRP7 target genes further emphasized its broad regulatory influence (Fig. 6f). However, investigating the correlation between GRP7 target gene binding efficiency and the change in mRNA TE under ABA treatment revealed no significant relationship (Fig. 6g, for detailed data process, see method). This suggests that GRP7 governs ABA signal-mediated mRNA TE independently of direct binding to those RNAs.

## Discussion

Plants, being sessile organisms, are constantly exposed to various abiotic stresses from their environment. To cope with these unfavourable conditions, plants have developed intricate strategies, including precise transcriptional response networks that sense and respond to harsh environmental cues, leading to redirection of their developmental programs. In addition to transcriptional changes, translation regulation represents another crucial pathway that contributes to the growth plasticity of plants. Despite its significance, the signaling mechanism responsible for linking abiotic stress and the protein translation machinery remains largely unknown. In this study, we focus on ABA signaling as an example to address this challenge.

ABA has long been recognized as a key phytohormone involved in various aspects of plant growth and stress responses^1,3,4^. Our investigation has uncovered that ABA treatment not only governs gene expression through transcription regulation but also significantly impacts the mRNA translation efficiency (Fig. 1). Specifically, ABA signaling inhibits the minor pre-rRNA processing pathway, disrupting the delicate balance of pre-rRNA processing. This imbalance leads to abnormal accumulation or reduction of rRNA maturation intermediates, thereby affecting ribosomal functions^8,9^. Moreover, ABA treatment induces alterations in the ribosome profiling pattern, influencing the global protein translation (Fig. 1). These translational changes are dependent on the core ABA signaling pathway involving receptors PYR/PYL/RCARs, the key protein phosphatase PP2Cs and kinase SnRKs (Fig. 2). Interestingly, while ABA signaling regulates transcriptional responses through important transcription factors such as ABI3/4/5, it also simultaneously regulates translation efficiency through specific RNA-binding proteins. We have identified highly expressed members of the RNA-binding protein family, GRP7, as crucial mediators for ABA’s regulation of global protein translation efficiency (Fig. 3, 4).

ABA treatment leads to a reduction in the protein levels of GRP7, a process that requires signaling transduction from receptor PYR/PYL/RCARs to key kinase SnRK2s. Thus, ABA signaling plays a dual role in regulating plant development, operating at both transcriptional and translational levels, possibly via distinct signaling pathways downstream of SnRK2s (Fig. 7). In the regulation of translation efficiency, the SnRK2-regulated protein levels of GRP7 appear to play a more predominant role.

**Fig. 7.**
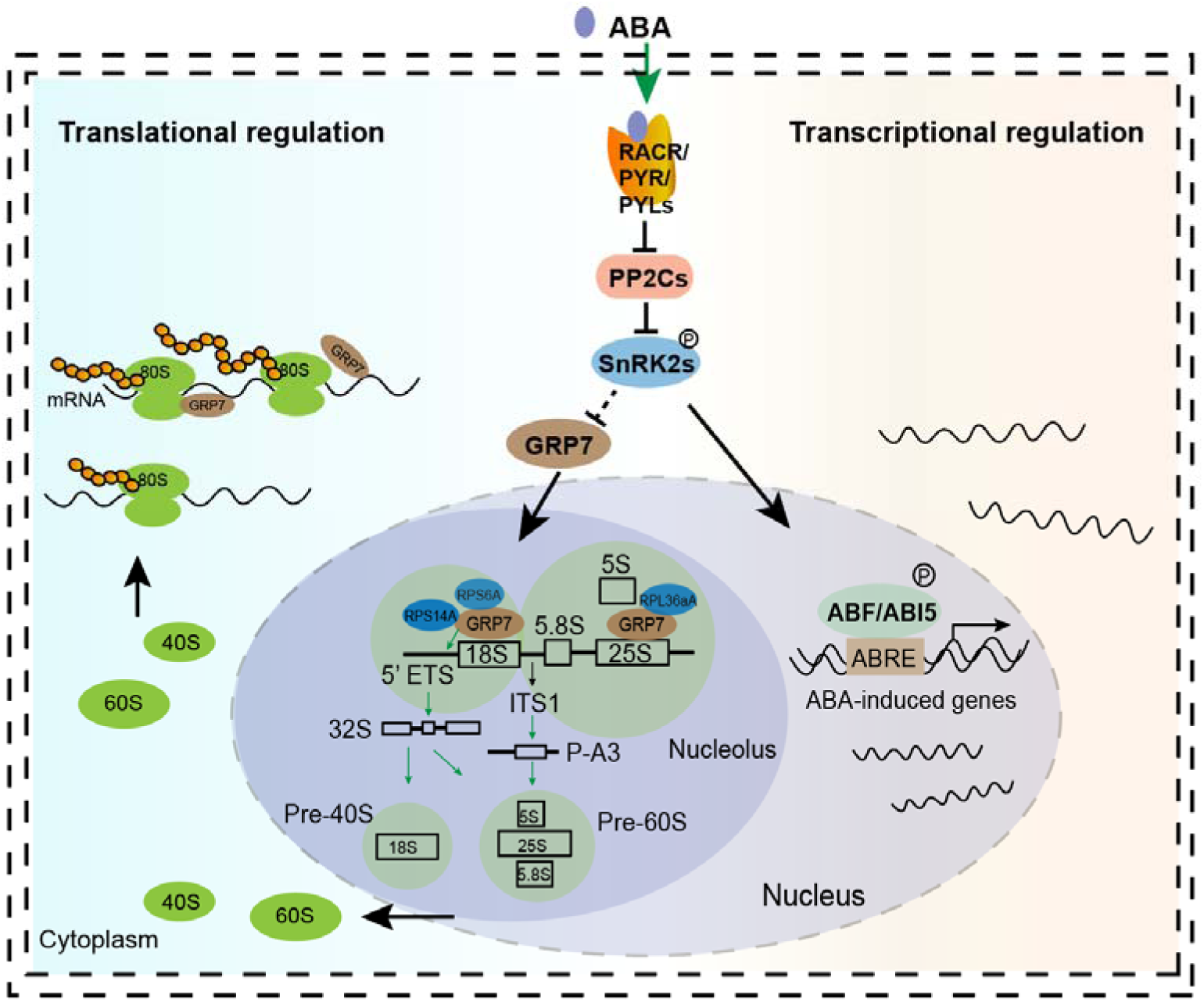
Working model for the regulatory role of GRP7 in ABA signal-mediated mRNA translation efficiency regulation. In addition to altering plant development through transcriptional regulation, ABA signaling also adjusts global mRNA translation efficiency. ABA represses GRP7 protein levels through its core signaling pathway factors RACR/PYR/PYLs, PP2Cs and SnRK2s, thereby inhibiting the 5’ETS first pre-rRNA processing pathway in the nucleolus by influencing GRP7 binding to pre-rRNA and interacting with ribosomal proteins. GRP7 actively participates in the protein translation process in the cytoplasm by its association with mature ribosomes.

Fascinatingly, the modulation of GRP7 protein levels by ABA signaling appears to be primarily achieved through alterations in its translation efficiency (TE), rather than through changes in transcription or protein stability. Investigating the specific components of the ABA signaling pathway responsible for orchestrating this effect warrants further in-depth analysis.

GRP7&8 are crucial regulators of multiple biological processes, encompassing mRNA splicing, primary microRNA processing, mRNA stability, and RNA chaperone activity, facilitating mRNA transport from the nucleus to the cytoplasm^52,56,57^. Our investigation sheds light on the involvement of GRP7&8 in translational regulation. Supporting their significant role in pre-rRNA processing, northern blot data revealed reduced levels of 32S rRNA in the *grp7grp8*^CR^ mutants. Additionally, polysome profiling data demonstrated decreased polysome levels and increased 80S monosome levels in the *grp7grp8*^CR^ mutants, indicating the influence of GRP7&8 on protein translation. Mechanistically, GRP7 directly binds to pre-rRNA in the nucleolus and physically interacts with ribosomal proteins RPS6A, RPS14S, and RPL36aA, promoting minor pre-rRNA processing (Fig. 7). In the cytoplasm, GRP7 co-fractionate with 40S, 60S, monosome, and polysomes, promoting protein translation efficiency (Fig. 7). Of note, GPR7 mediated mRNA translation efficiency change is likely not relied on the directly binding of GRP7 to the target mRNAs (Fig. 6). ABA treatment inhibits the accumulation of 32S rRNA, a marker of minor pre-rRNA processing, in Col-0 but not in the *grp7grp8*^CR^ mutant, suggesting that ABA might inhibit minor pre-rRNA processing partly through its effect on the protein level of GRP7. Similarly, ABA inhibits protein translational efficiency in Col-0 but only slightly in the *grp7grp8*^CR^ mutant, reinforcing the idea that ABA may influence mRNA translational efficiency partly through its effect on GRP7 protein levels. In conclusion, GRP7 play multifaceted roles in translation regulation and provide a novel molecular link between ABA signaling and the regulation of protein translation.

As GRP7 exerts influence on post-transcriptional processes of mRNA, including splicing and mRNA transport through direct binding^53^, its regulation of mRNA TE appears to operate independently of direct binding to target RNAs (Fig. 6). The intriguing question of how these distinct functions are separately executed warrants further exploration. One avenue for investigation could focus on the temporal response window to environmental stresses, along with the compartmentalization of its sub-cellular and sub-nuclear localization, as well as post-translational modification status. Additionally, considering that various abiotic stresses, such as drought and salinity, can trigger ABA biosynthesis and alter plant development^3,4^, delving into the role of the ABA-GRP7 module in mediating a broad spectrum of abiotic stress responses in plants would be of particular interest.

## Methods

### Plant materials and growth conditions

In this study, *Arabidopsis thaliana* plants used in this study were in the Col-0 background. The seeds were sown on 1/2 MS medium containing 1% (w/v) sucrose and 0.8% (w/v) agar, incubated at 4°C for 3 d, and transferred to a light incubator under a 16-h-light/8-h-dark cycle at 22°C.

To generate *pACTIN2::FLAG-RPL18,* the *RPL18* CSD sequence was amplified by PCR using cDNA as a template. The resulting DNA fragment was ligated into the pCAMBIA3301 vector driven by the *ACTIN2* promoter from *Arabidopsis thaliana*. The resulting construct was transformed into Agrobacterium tumefaciens GV3101 and transferred to Col-0 plants using the floral dip method. The *grp7grp8*^CR^-A, *grp7grp8*^CR^-G, *grp8*^CR^-A and *grp8*^CR^-G mutants were created by CRISPR/Cas9 editing. *grp7-1*^45^ (SALK_039556) mutant was described previously. *pp2c* triple mutant (*3m*) (*abi1-2abi2-2hab1-1*)^5^, *snrk2.2/2.3/2.6*^43^, *nced6*^58^ and *pyr1pyl1/2/4*^42^ was described previously. *pGRP7::GRP7-GFP grp7-1* complementary line^53^ was described previously. The primers used for identification of the mutations are listed in Supplementary Data 10.

### Green cotyledons assay

For the green cotyledons assay, ∼50 seeds were sown on 1/2 MS medium containing different concentrations of ABA. The plates were transferred from a temperature of 4°C to the light incubator, where they were grown for 15 d to examine the seed germination ratio, with three plates prepared per experiment. Seedlings with expanded cotyledon were considered as green cotyledons.

### Northern blot analysis of pre-rRNA processing

Northern blot assays were conducted as previously described^9^ with some modifications. Total RNA was isolated using RNA extraction kit according to the manufacturer’s instructions. Five micrograms of total RNA was separated on a 1.4% (wt/vol) agarose/formaldehyde gel and transferred to a Hybond N+ membrane. For short DNA probes, oligonucleotides labeled with biotin by Biotin 3’ DNA labeling kit (Thermo Fisher, 89818) were used to detect precursor rRNA. Hybridization was performed overnight at 45°C in Church buffer. The blots were washed and applied to chemiluminescent nucleic acid detection module (Thermo Fisher, 89880). then signals were detected by Chemiluminescence imaging. The probes are listed in Supplementary Data 10.

### Polysome profiling analysis

*Arabidopsis* polysomes were fractionated over sucrose gradients as described^59^ with minor modifications. In brief, 3-day-old seedlings were treated with 5 μM ABA for 4 h and then ground in liquid nitrogen followed by resuspension in polysome extraction buffer. Supernatant was loaded onto a 10%–50% sucrose gradient and spun in a Beckman SW41Ti rotor at 33,500 rpm for 4 hr at 4°C. We collected 12 fractions by a gradient fractionator.

### RNA immunoprecipitation (RIP) and qPCR analyses

One gram of 3-day-old seedlings were treated with 5 μM ABA for 4 hr and then ground in liquid nitrogen followed by crosslinking twice at 600 mJ/cm^2^ in a UVP crosslinker (Analytik jena). The nuclei were extracted as previously described^60^. The RIP experiment was performed as described by^61^. The protein A beads bind GRP7 antibody were used for IP. Finally, the immunoprecipitated RNA was reverse transcribed with random primers (TransGen, AH301-02). Gene-specific primers were used for RT–PCR to detect the target transcripts as described^55^.

### Ribosome profiling (Ribo-seq) experiment

Ribosome profiling experiment was performed as previous reported with some modifications^62^. In brief, 3-day-old seedlings were treated with 5 μM ABA for 4 h and then ground in liquid nitrogen followed by resuspension in 600 ml ice-cold lysis buffer. Clarifying the lysate by centrifugation for 10 min at 20,000 g under 4°C, the soluble supernatant was recovered. 6 ml of RNase I (100 U/ml) was added to 600 ml lysate and Incubated for 45 min at room temperature with gentle mixing. 10 ml SUPERase-In (Invitrogen Cat# AM2694) RNase inhibitor was then added to stop nuclease digestion. Meanwhile, MicroSpin S-400 HR columns (GE Healthcare Cat# 275140-01) were equilibrated with 3 ml of mammalian polysome buffer by gravity flow and emptied by centrifugation at 600 g for 4 min. Then digested lysate was immediately loaded on the column and eluted from the column by centrifugation at 600 g for 2 min. The RNA was extracted from the flow-through using Trizol (Thermo Fisher, 15596018CN). The ribosomal RNA fragments were removed using the Ribo-off rRNA depletion kit (Plant) (Vazyme, N409) and separated on a 15% denaturing urea-PAGE gel. The size ranges from 27 nt to 30 nt was cut and thus obtained RNA fragments were subjected into library generation using Smarter smRNA-Seq kit (Takara Cat# 635031).

### Crosslinking and Immunoprecipitation followed by sequencing (CLIP-seq)

CLIP-seq was performed as described previously^54^ with some modifications. 3-day-old seedlings were treated with 5 μM ABA for 4 h and then ground in liquid nitrogen followed by crosslinking twice at 600 mJ/cm^2^ in a UVP crosslinker (Analytik jena). The RNA-protein complexes (RNP) were enriched from the lysate by immunoprecipitation using GFP-Trap (Lablead, GNM-25-1000) and were partially digested by micrococcal nuclease (2 × 10^−5^ U/μL, Takara, 2910A). The digested RNA was ligated to the 3′-RNA adaptor labeled by biotin. The RNPs were separated by SDS-PAGE and were further transferred to nitrocellulose membrane. Membrane pieces corresponding to the RNP-containing were selected to recover RNA fragments. RNPs were treated with Proteinase K, and reverse transcription was performed. The purified cDNA was ligated to the 5′-DNA adaptor.

### IP-MS assay

Immunoprecipitation of GRP7-GFP proteins from 3-day-old seedlings with or without ABA treatment was performed as previously described^63^ with some modifications. Proteins were solubilized from plant powder using buffer A (25 mM Tris-HCl pH 7.5, 150 mM NaCl, 15 mM MgCl_2_, 1 mM DTT, 8% glycerol, 1% Triton X-100, 0.5% sodium deoxycholate, 100 mg/ml Cycloheximide (CHX), 100 U/ml SUPERase In RNase Inhibitor, 25 U/ml Turbo DNase, Protease Inhibitor). GFP-Trap (Lablead, GNM-25-1000) were used to immunoprecipitate GRP7-GFP from the protein extracts. IP samples were first washed 3 times each for 5 min at 4°C with buffer B (25 mM Tris-HCl pH 7.5, 150 mM NaCl, 15 mM MgCl_2_, 1 mM DTT, 1% Triton X-100, 0.5% sodium deoxycholate, 100 mg/ml CHX). Afterward, beads were washed 3 times for 5 min each at 4°C using buffer C (25 mM Tris-HCl pH 7.5, 300 mM NaCl, 15 mM MgCl_2_, 1 mM DTT, 1% Triton X-100, 0.5% sodium deoxycholate, 100 mg/ml CHX).

### Yeast two hybrid assay

The coding sequences of GRP7 was fused in-frame with the GAL4 DNA binding domain (BD) of the bait vector pGBKT7. The coding sequence of RPS6A, RPS14A and RPL36aA were cloned in the prey vector pGADT7. The bait plasmid and the prey plasmid were co-transformed into the yeast strain AH109. The clones growing in the –L/-T liquid selection medium (Clontech) were diluted by a 10x series dilution and spotted onto –L/-T/-H medium (Clontech) to determine interactions between the two proteins.

### BiFC assay

The coding sequences of GRP7 were cloned into the pSCYNE(R) vector, and RPS6A, RPS14A and RPL36aA were cloned into the pSCYCE(R) vector. The plasmids were then transformed into Agrobacterium GV3101 cells, and different combinations were mixed for injection into *Nicotiana benthamiana* leaves with the silencing suppressor P19 strain and nucleolus maker FIB2-mCherry. After 3 days, CFP signals were observed under a confocal laser-scanning microscope (Zeiss).

### Western blot assay

For western blotting analysis, protein samples were separated on SDS-PAGE and transferred onto PVDF membrane. After blocking with 5% milk in PBST (0.1% Tween-20) for 1 h, the membrane was incubated with anti-GRP7^45^, anti-FLAG (Sigma, F1804), anti-RPS6 (PHYTOAB, PHY2025A) for 1 h at room temperature. Protein levels were detected by Tanon-5200 system (Tanon).

### Subcellular localization analysis

For subcellular localization analysis of GFP and GRP7-GFP fusion protein, 1.4 kb of the GRP7 promoter and the GRP7 5’UTR, intron and coding sequences was fused with GFP, transiently expressed in *Nicotiana benthamiana* leaf epidermal cells with the silencing suppressor P19 strain and nucleolus maker FIB2-mCherry. After 2 days, cells expressing GFP or GRP7-GFP fusion proteins were observed and photographed by confocal laser-scanning microscope (Zeiss).

### Bioinformatics analysis

Raw reads of RNA-seq, Ribo-seq, and CLIP-seq were filtered by fastp v0.20.1 for adapters removing, low-quality bases trimming, and reads filtering^64^. RNA-seq reads were filtered with parameter “--detect_adapter_for_pe”, CLIP-seq reads were filtered with parameter “--adapter_sequence TGGAATTCTCGG --adapter_sequence_r2 GATCGTCGGACT”. For Ribo-seq data, only first sequencing reads (*_R1.fq.gz) were used and filtered with parameter “-a AAAAAAAAAA -f 3 -l 16”. The high-quality reads of RNA-seq were mapped to the Arabidopsis thaliana genome (TAIR10) using STAR (v2.7.10) with default parameters^65^. The clean reads of Ribo-seq were firstly aligned against the non-coding RNA sequences of *A. thaliana* downloaded from Ensembl Plants^66^ using bowtie2^67^ to produce the unaligned reads. The unaligned reads were mapped to reference genome using STAR with parameters “-- outFilterMismatchNmax 2 --outFilterMultimapNmax 1 --outFilterMatchNmin 14 -- alignEndsType EndToEnd”. For CLIP-seq data, the clean reads were mapped to reference genome using STAR with no more than two mismatches^65^ and the duplicates in mapped reads were removed using Picard v2.23.3.

Two or three replicates bam files were merged using Samtools v1.4^68^. To normalize and visualize the individual and merged replicate datasets, the BAM files were converted to bigwig files using bamCoverage provided by deepTools v3.3.0 with 1 bp bin size and normalized by RPKM (Reads Per Kilobase per Million mapped reads) with parameters “-bs 1 --effectiveGenomeSize 120,000,000 --normalizeUsing RPKM --smoothLength 5”^69^. The number of reads that mapped to each gene was counted using featureCounts v2.0.1 with the parameter “-p -P -B -C” for RNA-seq and default parameters for Ribo-seq^70^. The raw counts were further normalized to FPKM (Fragments per kilobase per million mapped reads) for RNA-seq and RPKM (Reads per kilobase per million mapped reads) for Ribo-seq. FPKM or RPKM values of genes were Z-scaled and clustered by k-means method and displayed using R package ComplexHeatmap (v2.4.3)^71^. Gene Ontology enrichment was performed using an R package clusterProfiler v3.18.1^72^.

The raw counts files of RNA-seq or Ribo-seq were used as inputs for differentially expression analysis by DESeq2 v1.26.0^73^. Genes with > 1 RPKM in ribo-seq and > 1 FPKM in RNA-seq were kept for the calculation of translation efficiency (TE). The TE of a gene was calculated as the ratio between the average RPKM values in the Ribo-seq and FPKM values in the RNA-seq. The formula is RPKM Ribo-seq/FPKM RNA-seq. The raw counts matrix of RNA-seq and Ribo-seq were implemented into Xtail package (v1.1.5) to calculate the differential translation efficiencies^74^. The threshold for differential translational efficiency was *p* value.adjust L<L0.05 (Supplementary Data 1-4, 6-7). The slope and intercept were computed use standard major axis (SMA) method in R package “lmodel2” (Fig. 4b). Comprehensive gene expression analysis in *grp7grp8*^CR^-A mutant in comparison with that in Col-0 was showed that the transcription and TE does not up/down-regulated in *grp7grp8*^CR^- A mutant among the ABA-induced transcriptionally altered and TE-altered genes in Col-0 (Fig. 4d).

The GRP7 binding sites were identified by performing CLIP-seq peak calling using CLIPper (v0.1.4) with default parameters^75^. Then, bedtools was used to retain the peaks that were identified in both biological replicates^76^. Finally, we defined the peaks that were only identified in IP samples but depleted in negative controls as significant binding sites of GRP7. The GRP7 binding peaks were annotated to the Arabidopsis thaliana genome using the “annotatePeak” function in R package ChIPseeker (v1.26.2)^72^. The gene promoters are defined as 1.5 Kb upstream of gene TSS. The GRP7-binding efficiency of a gene was calculated as the ratio between the CPM values (Counts per million) in the CLIP-seq and the FPKM values in the mRNA-seq (Fig. 6g). To investigate motifs enriched in GRP7 binding sites, HOMER was used to identify de novo motifs using the command “findMotifsGenome.pl <foreground> tair10 <output location> -rna -p 4 -bg <background>”. The foreground was a bed file of significant peaks; the background was randomly defined peaks within the same annotated region as the foreground peaks (Supplementary Data 8).

### Statistics and data visualization

R (https://cran.r-project.org/;version 4.0.2) was used to compute statistics and generate plots if not specified. The Integrative Genomics Viewer (IGV) was used for the visual exploration of genomic data^77^. The correlation between two groups of data was conducted with the Pearson analysis (Fig. 1c, 6g, Fig. S1c) and *P* values are determined by the two-sided Pearson correlation coefficient analysis (Fig. S1b). For enrichment analysis, Hypergeometric test (Fig. 4e, 5d, 6f, Fig S1e, S6c) were used. For two groups’ comparison of data, the student’s *t*-test was used, such as Fig.1f, 1g, 1i, 2b, 2c, 2d, 2f, 2g, 3c, 3d, 3f, 4f, 5c, 6c, Fig. S3b, S3c, S3e, S4c, S5a, S5b, S6b.

### Data availability

The raw sequence data of RNA-seq, Ribo-seq, and CLIP-seq in this study were deposited in the Genome Sequence Archive (https://bigd.big.ac.cn/gsa)^78^ in National Genomics Data Center^79^ under accession number CRA012798.

### Code availability

Code used for all processing and analysis is available at Github (https://github.com/yx-xu/GRP7-mediate-translational-regulation).

## Author contributions

J.X. designed and supervised the research, J.X., J.Z. and Y.-X.X. wrote the manuscript. J.Z. did most of the experiments; Y.-X. X. performed bio-informatics analysis; F.-A. T. constructed *pACTIN2::FLAG-RPL18* and help with polysome profiling assay; C.-S.H. help with CLIP-seq; Y.-L.D., and X.-F.Y. provided *pGRP7::GRP7-GFP grp7-1* complementary line and put forward valuable advice on the research; C.-S.H, X.Y., X.-F. Y. and Y.-L.D. revised the manuscript; J.Z., Y.-X.X. and J.X. prepared all the figures. All authors discussed the results and commented on the manuscript.

## Acknowledgements

We thank Professor Z.-Z.Gong (CAU) provided *pp2c 3m*, Professor J.-G.Huang (SDAU) and Professor J-K Zhu (SUSTC) provided *snrk2.2/2.3/2.6* seeds, Professor Q.Xie (IGDB, CAS) provides *nced6* and *pyr1pyl1/2/4* seeds. We thank Professor W.F. Qian (IGDB, CAS) for supporting ribo-seq analysis, Professor J. S. Zhang (IGDB, CAS) for suggestions of CLIP-seq analysis and Dr. C.Y. Liu (IGDB, CAS) for help with Northern blot assay for pre-rRNA processing. We thank Drs. B.Y. Shi (IoG, CAS) and J. Wang (IGDB, CAS) for suggestions on the experimental conduction and data analysis of Ribo-seq assay.

---

**Supplementary Fig. 1.**
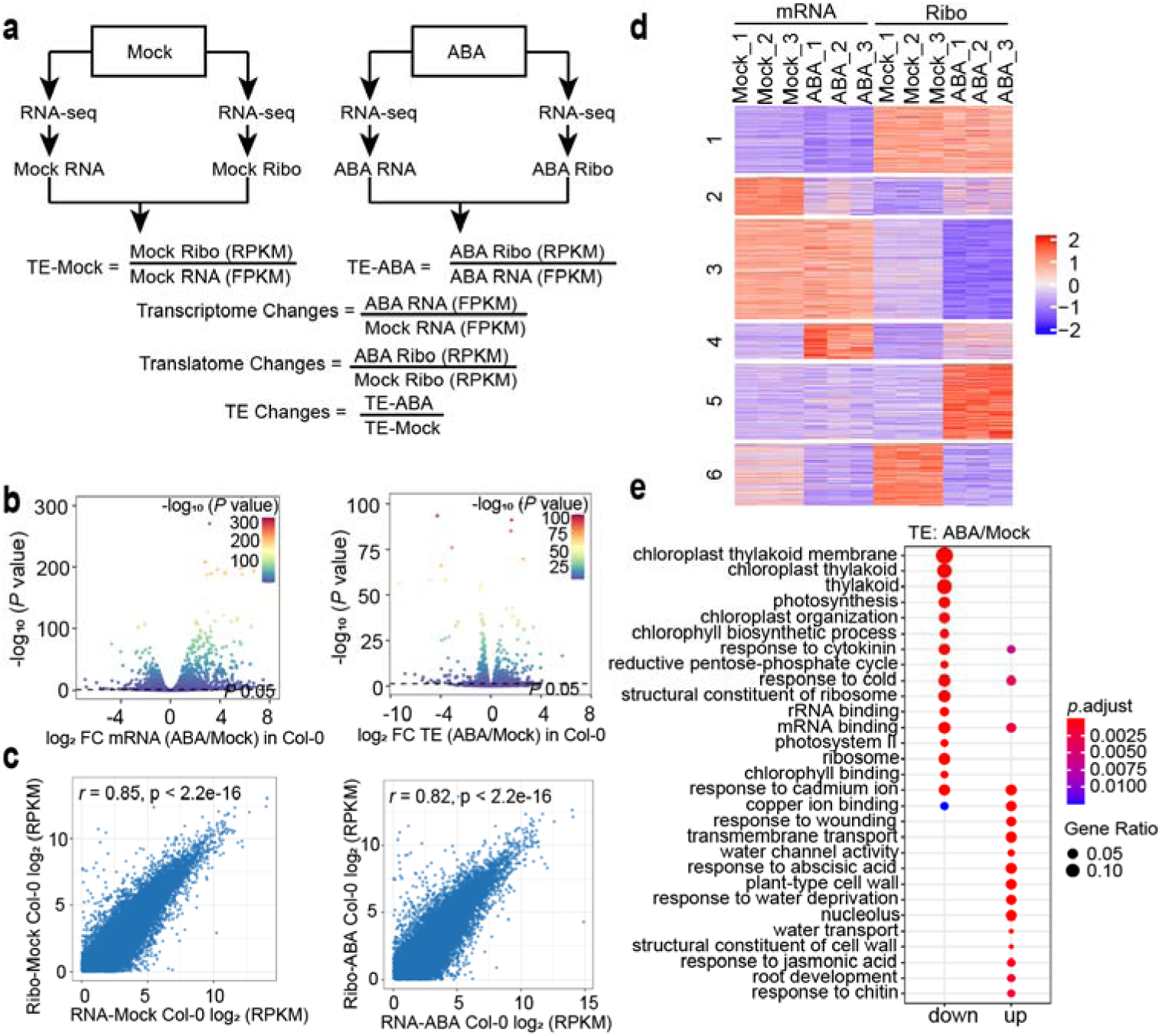
Overview analysis of the RNAseq and Ribo-seq in Col-0 under mock and ABA conditions. **a** Schematic illustrating the data processing workflow for analyzing transcriptional and translational changes in Col-0 upon ABA treatment. **b** Analysis of ABA-induced alterations at the transcription and translation efficiency (TE) levels in Col-0. The black dashed line indicates the 0.05 *P* value cutoff. **c** Pearson correlation coefficient between RNA-seq and Ribo-seq. **d** Heatmap displaying transcription and translation changes of genes with altered TE in Col-0 after ABA treatment. **e** GO functional categories of genes with up- and down-regulated TE change in Col-0 after ABA treatment.

**Supplementary Fig. 2.**
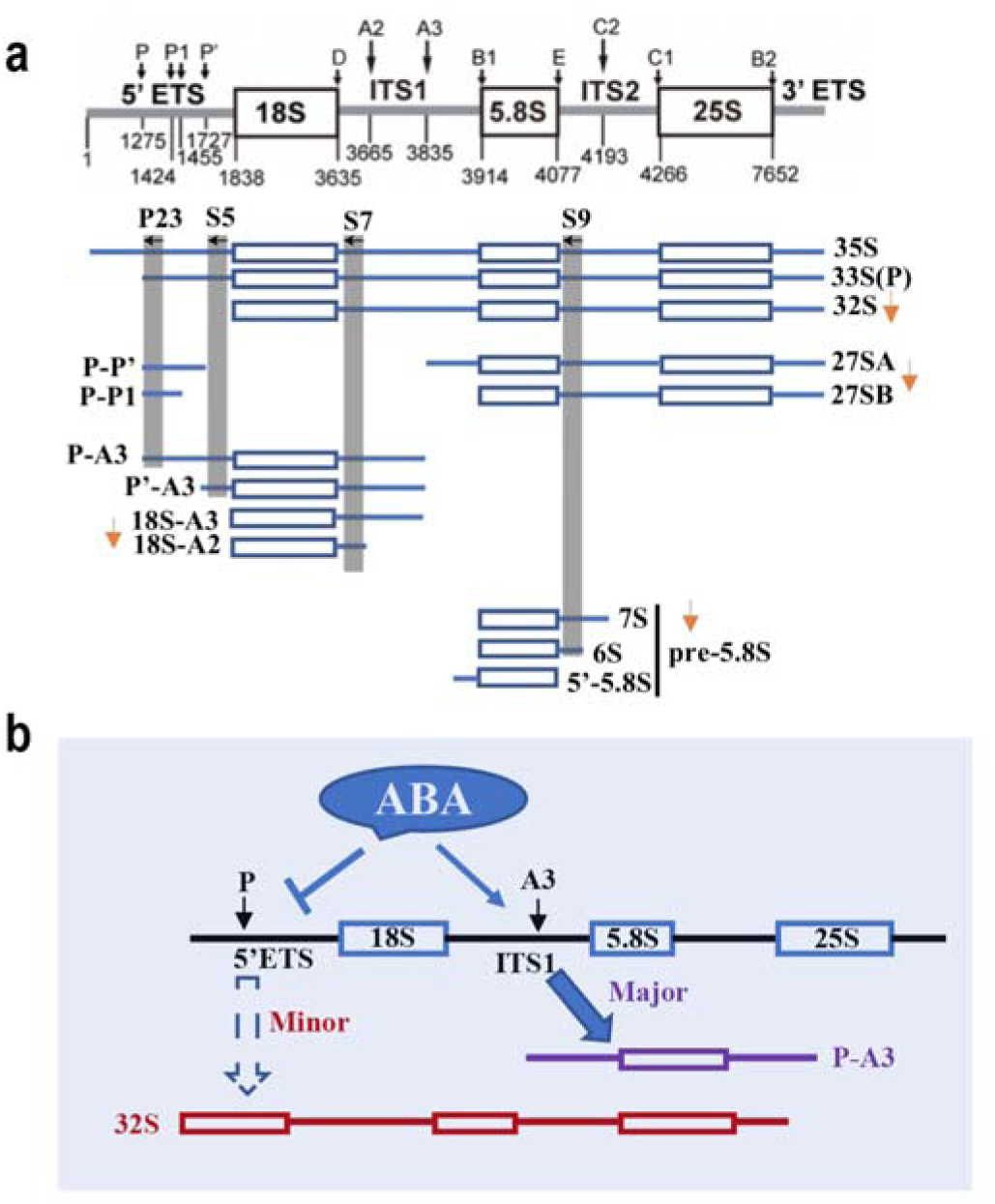
ABA influences pre-rRNA processing. **a** Diagram illustrating various pre-rRNA processing intermediates detected by Northern blots with specific probes, indicated by horizontal arrows. Orange arrows highlight decreased rRNA intermediates by ABA treatment. **b** Working model for the influence of ABA in pre-rRNA processing. Two pre-rRNA processing pathways, different in cleavage order but not cleavage site, coexist in Arabidopsis. The P-A3 fragment and 32S rRNA are diagnostic for the major and minor pathways, respectively. ABA causes unbalanced pre-rRNA processing pathways with up-regulation of the major pathway (P-A3) and down-regulation of the minor pathway (32S rRNA), leading to pre-rRNA processing defects.

**Supplementary Fig. 3.**
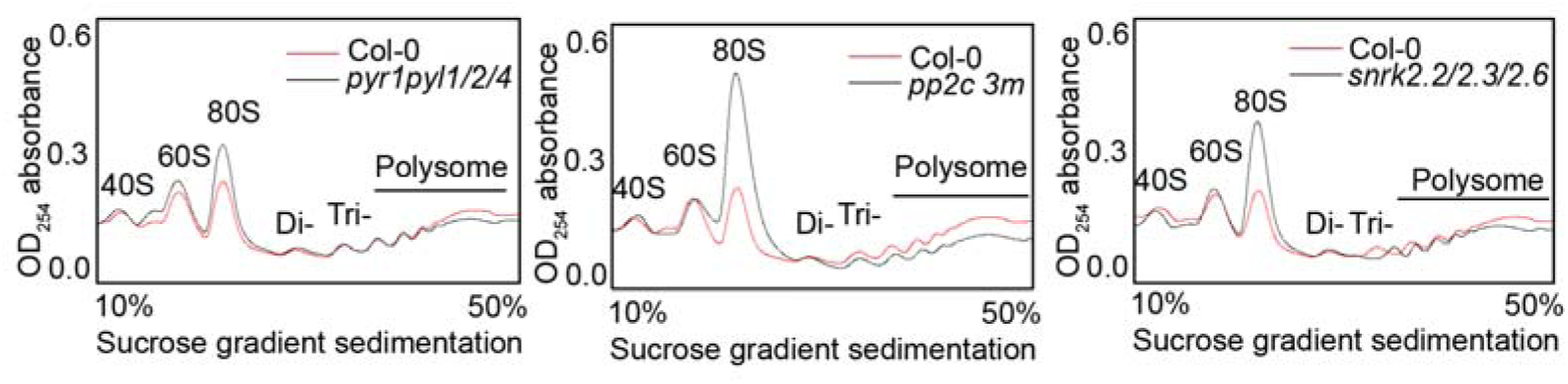
Polysome profiles of ABA signaling pathway factor mutants under mock condition. Polysome profiles were analyzed by sucrose gradient sedimentation, and OD_254_ was measured for the Col-0 and ABA signaling pathway factor mutants, *pyr1pyl1/2/4*, *pp2c 3m* and *snrk2.2/2.3/2.6*.

**Supplementary Fig. 4.**
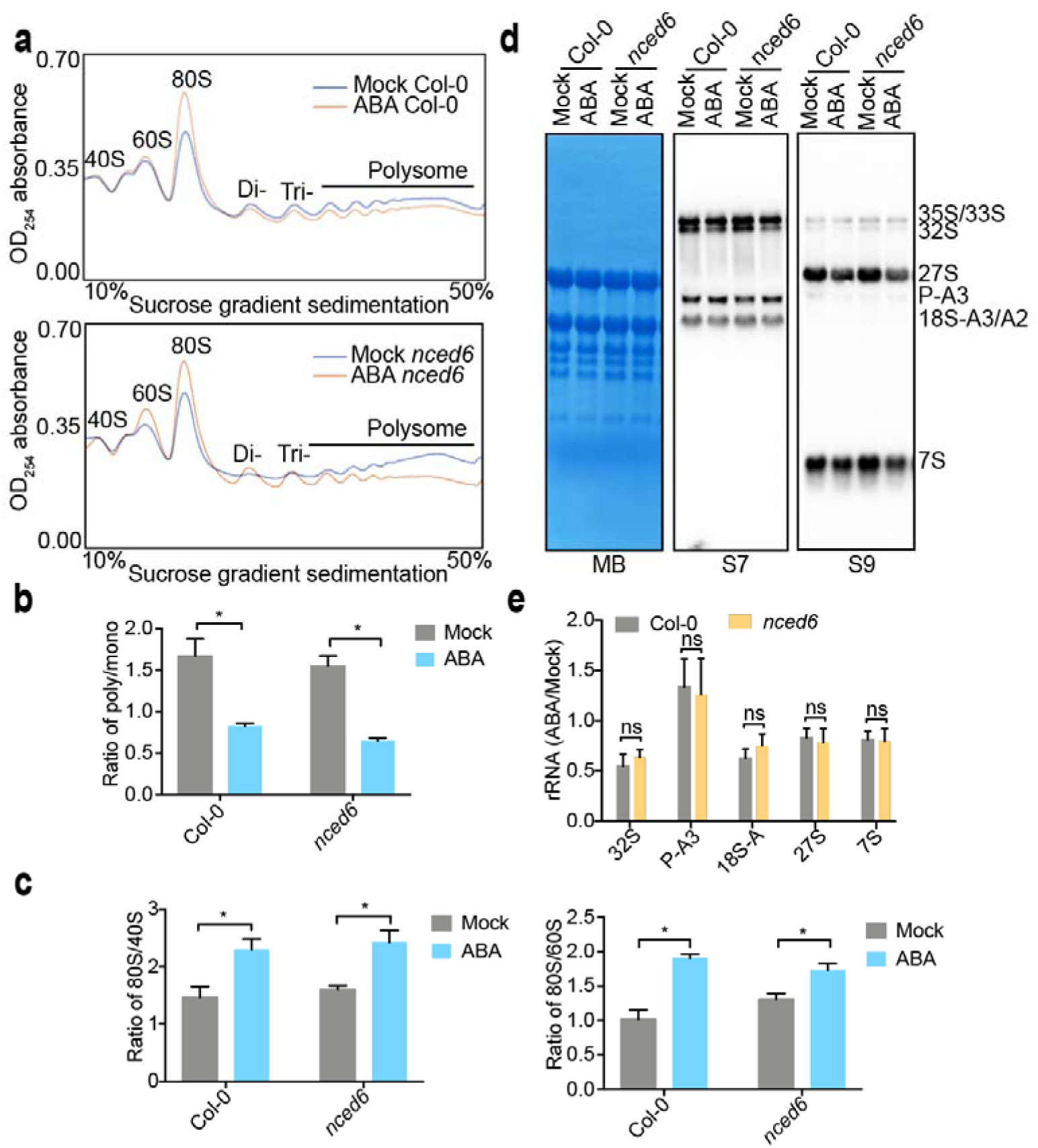
ABA influences translation efficiency independent of the biosynthesis enzyme NCED6. **a** Polysome profiles of Col-0 and *nced6* under mock and ABA conditions. **b** Quantification of polysome/monosome (poly/mono) ratios for data presented in **(a).** Student’s *t* test: \**p* < 0.05; \*\**p* < 0.01; \*\*\**p* < 0.001. Two bio-replicates were performed. **c** Ratios of 80S/40S and 80S/60S quantified for the data shown in **(a)**. Student’s *t* test: \*\**p* < 0.01. Two bio-replicates were performed. **d** Northern blot identification depicting the pre-rRNA processing in Col-0 and *nced6* with mock and ABA treatment. Methylene blue staining (MB) serves as loading control. **e** Quantification of the intensities of different rRNA intermediates. Values were normalized to methylene blue stain for mature 25S and 18S rRNAs and then expressed as a ratio to the intensity observed in Col-0 under mock treatment. The baseline is set to 1, and the standard deviation is indicated as error bar. Student’s *t* test: ns, no significance. Two bio-replicates were performed.

**Supplementary Fig. 5.**
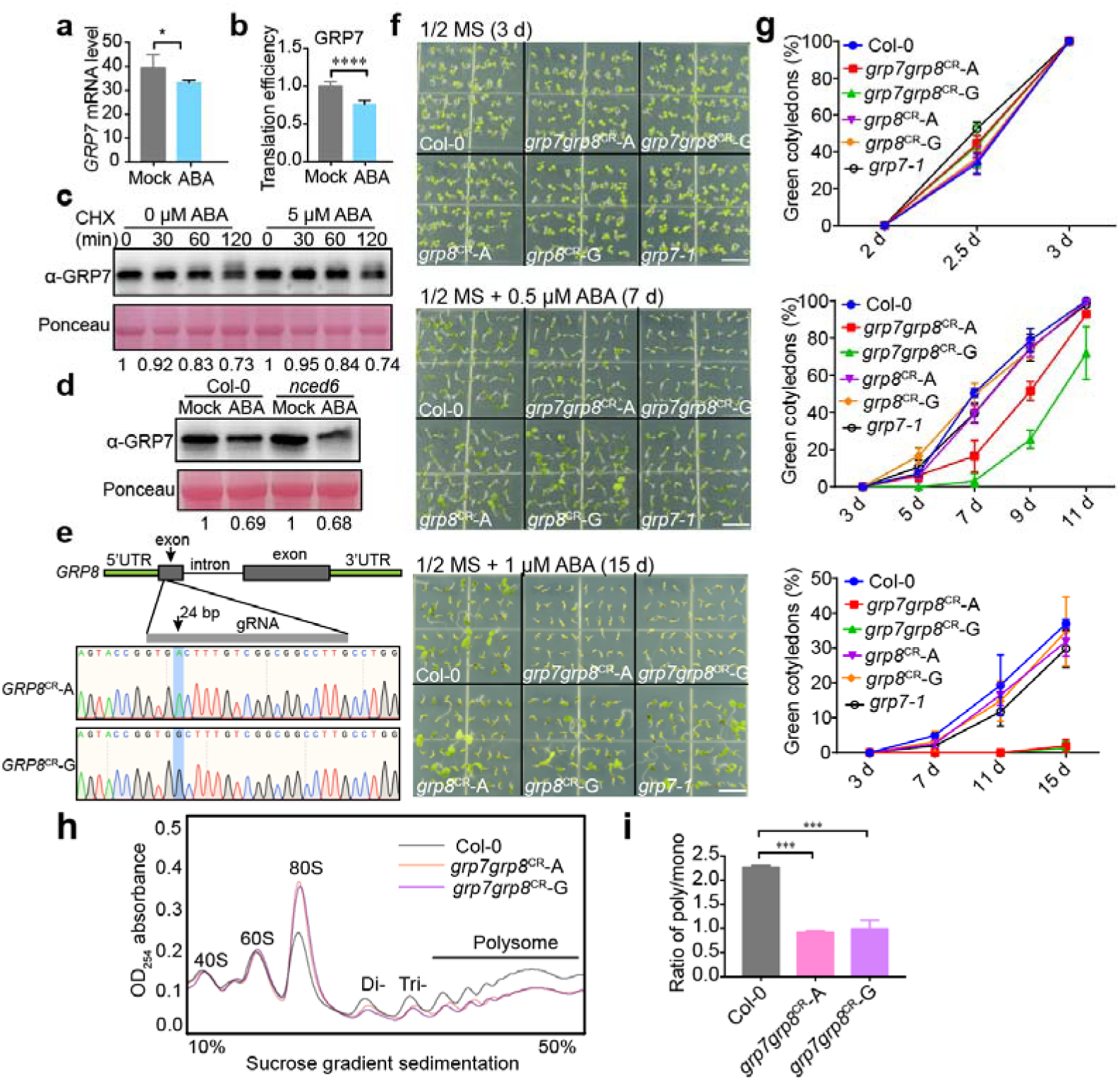
ABA influences GRP7 level and generation of *grp7grp8* double mutant. **a** Quantitative RT-PCR measured relative *GRP7* mRNA expression in total RNA from 3-day-old Col-0 seedlings treated with mock or ABA. Internal reference: *PP2A*. Student’s t-test: *p< 0.05. Three repeats were performed. **b** TE of the GRP7 mRNAs, calculated as their relative expression in polysomal/total RNA fractions, Student’s *t* test: \*\*\*\**p* < 0.0001. Three repeats were performed. **c** Degradation of GRP7 in protein extracts from Col-0 seedlings treated with 150 μM CHX (to block new translation), 5 mM ATP, and 0 or 5 μM ABA for indicated time points. **d** Protein levels of GRP7 in Col-0 and *nced6* after ABA treatment detected with anti-GRP7 antibody. **e** Construction of GRP8 CRISPR mutants in *grp7-1*, revealing single base A or G insertion at the gRNA targeting region. **f** Photographs of seedlings at 3 dps (days post-stratification) on 1/2 MS medium containing 0 μM ABA, at 7 dps on 1/2 MS medium containing 0.5 μM ABA, and at 15 dps on 1/2 MS medium containing 1 μM ABA. (Scale bars, 5 mm). **g** Percentage of seedlings showing green cotyledons in (**f**) analyzed. **h** Polysome profile analyzed by sucrose gradient sedimentation, with OD_254_ measured for Col-0 and *grp7grp8*^CR^*-*A/G lines. **i** Quantification of polysome/monosome (poly/mono) ratios for data presented in **(h).** Student’s *t* test: \*\*\**p* < 0.001. Two bio-replicates were performed.

**Supplementary Fig. 6.**
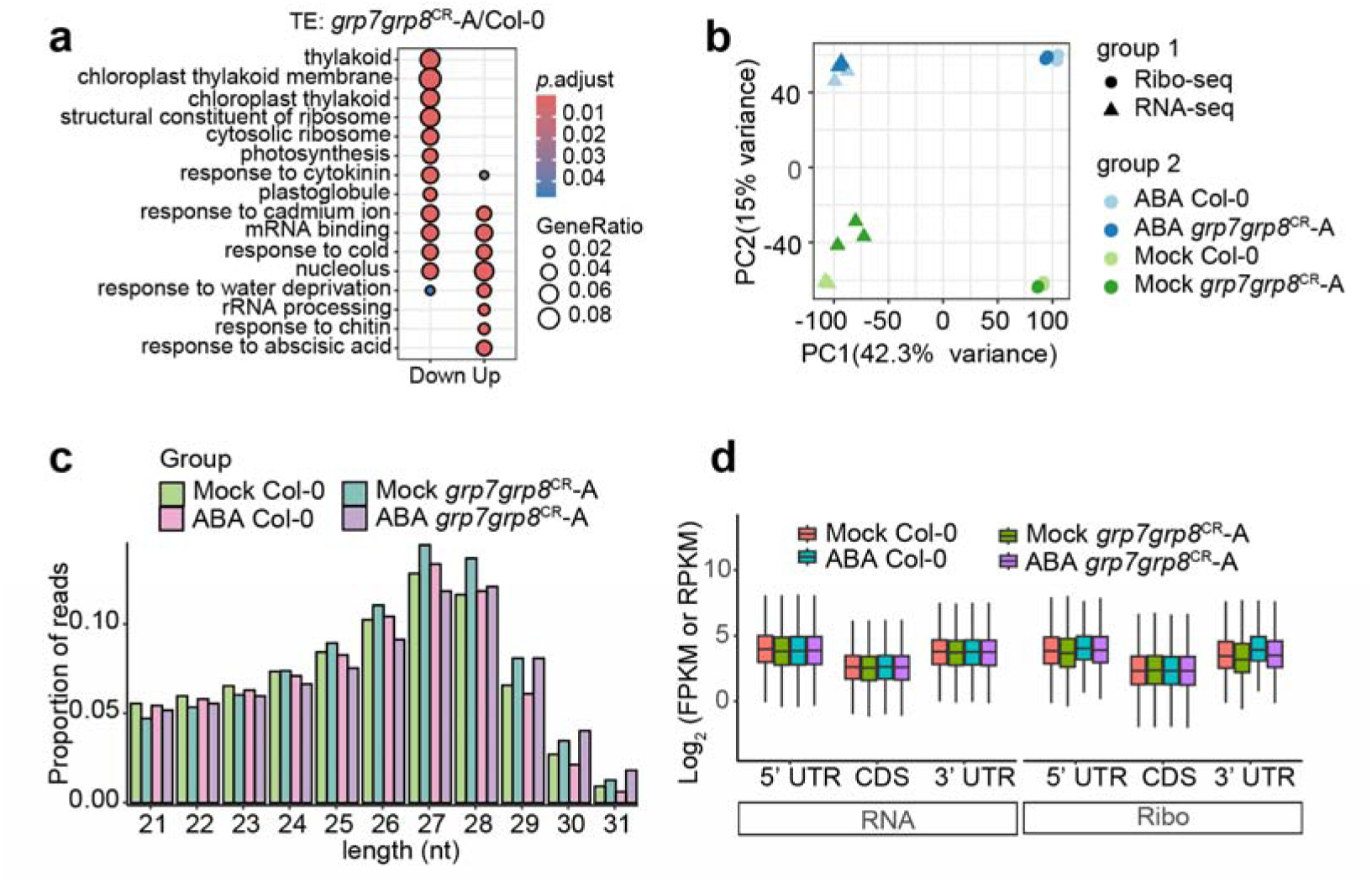
Quality control and reproducibility of RNA-seq and Ribo-seq libraries. **a** Biological process GO enrichment analysis of genes down- and up-regulated TE in *grp7grp8*^CR^-A. Fold enrichment in each GO category is depicted by the size of points, and the *p*.adjust value is indicated by the color scale. **b** Principal component analysis (PCA) of the RNA-seq and Ribo-seq from Col-0 and *grp7grp8*^CR^*-*A. **c** Length distribution of total reads from Ribo-seq libraries. **d** Read density along 5′ UTR, CDS, and 3′ UTR of total reads from RNA-seq and Ribo-seq libraries

**Supplementary Fig. 7.**
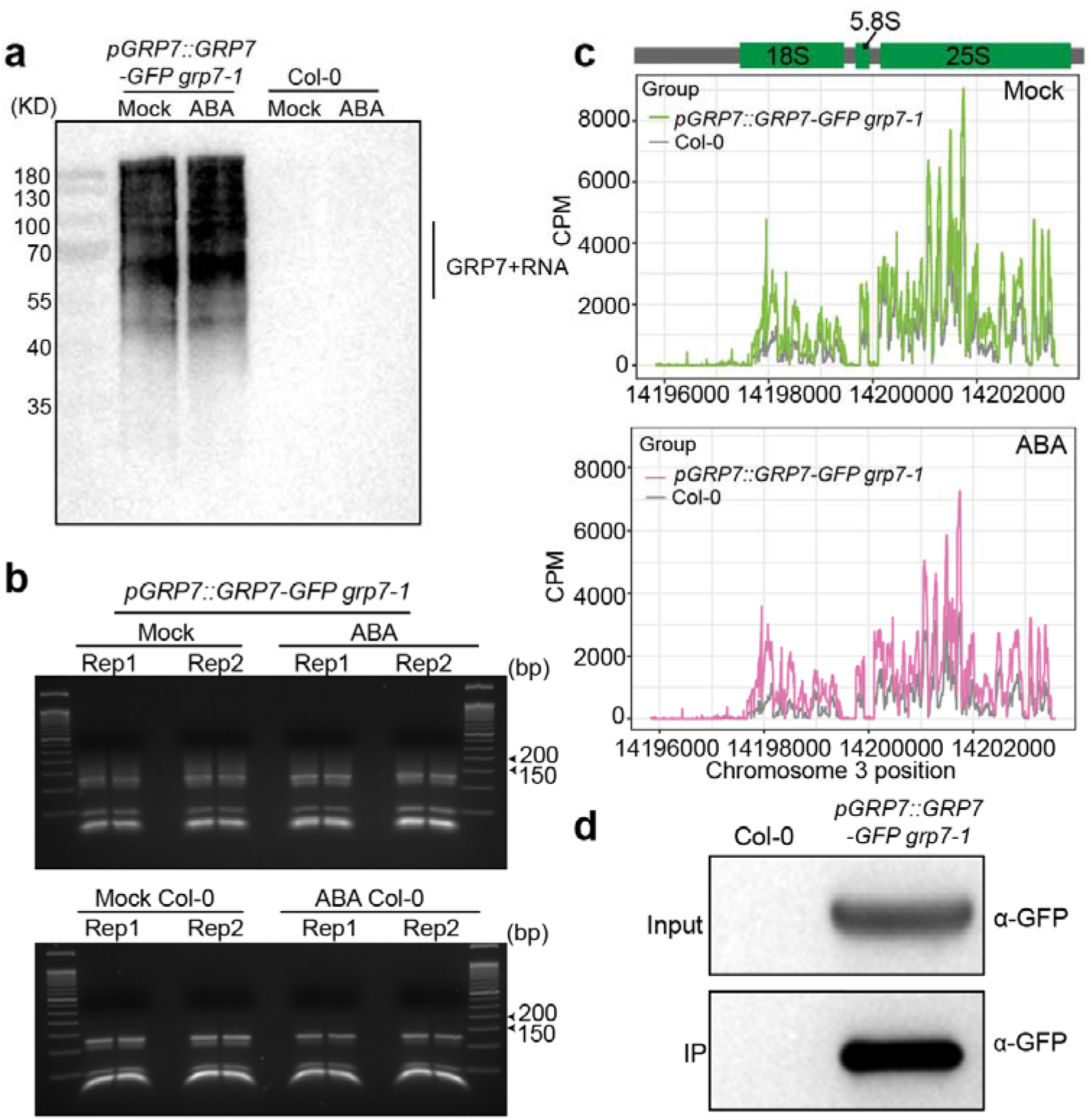
CLIP-seq reveals GRP7 binding to rRNA. **a** Three-day-old seedlings of complementary lines *pGRP7::GRP7-GFP grp7-1* and Col-0 after ABA treatment were utilized for CLIP-seq library construction. Immunoprecipitation (IP) was performed using GFP-Trap. The selected regions containing the GRP7-RNA complex are demarcated by a black line. **b** Size selection of the fragments’ length in the CLIP-seq libraries. The final PCR products with lengths ranging from 150 to 200 bp were size-selected. After PCR amplification, the libraries were separated by a 3% agarose gel. A 50 bp DNA ladder with the unit “bp” is labeled on the right. **c** Locations of GRP7 binding sites on rRNAs with mock (green) and ABA (pink) in chromosome 3, gray line serves as Col-0 control. **d** IP-MS for detecting the interacting proteins of GRP7 of three-day-old seedlings of *pGRP7::GRP7-GFP grp7-1* and Col-0. Col-0 was used as the negative control.

**Supplementary Fig. 8.**
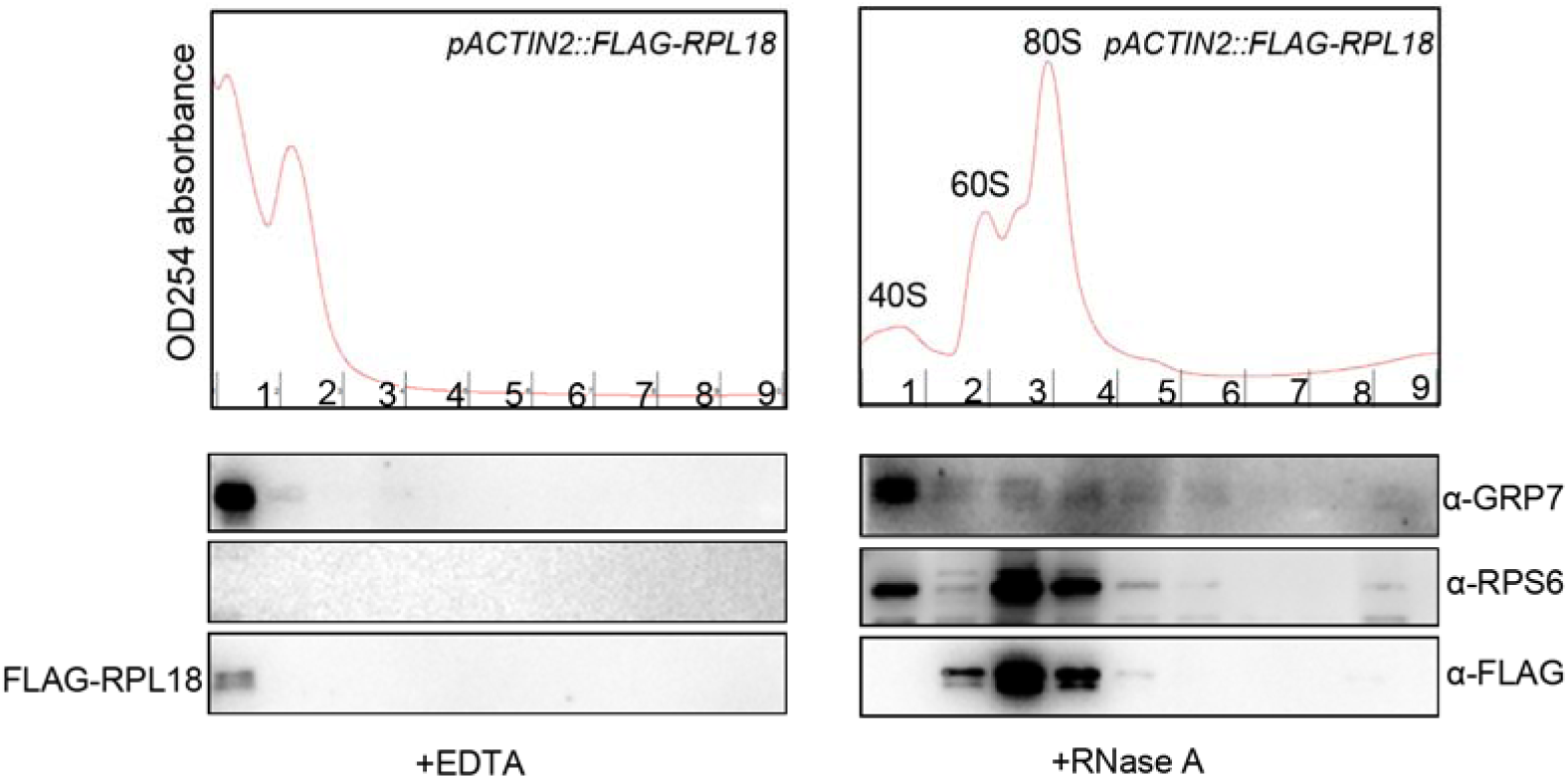
GRP7 associates with mature polysomes. Lysates with EDTA or RNase A treatment were subjected to sucrose gradient fractionation. Collected fractions were precipitated and analyzed for the presence of GRP7, RPS6, and FLAG-RPL18 by western blot using anti-GRP7, anti-RPS6, and anti-FLAG antibodies.

